# Force-Enhanced Sensitive Detection of New DNA-Interactive Agents from Microorganisms at the Single-Molecule Level

**DOI:** 10.1101/2024.04.22.590585

**Authors:** Tianyu Liu, Teng Cai, Hongwei Liu, Aiying Li, Meng Yin, Yan Mei, Yueyue Zhou, Sijun Fan, Yao Lu, Luosheng Wan, Huijuan You, Xiaofeng Cai

**Author notes:** These authors contributed equally to this work (Tianyu Liu, Teng Cai).

## Abstract

The discovery of microbial-derived DNA-interacting agents, which hold broad therapeutic potential, is inherently challenging due to the limited sensitivity and specificity of conventional methodologies. Our study introduces a pioneering application of single-molecule stretching assay (SMSA) in natural product chemistry to identify DNA-intercalating agents directly from microbial cultures or extracts. We demonstrate that mechanical force can enhance sensitivity by increasing both the binding affinity *K*a and the quantity of ligands bound. The changes induced by intercalators in the counter length and overstretching transition of dsDNA yield a distinctive and highly specific signature indicative of DNA intercalative binding, thereby enabling straightforward detection of DNA intercalators even in trace amounts from microbial cultures. This methodology eliminates the need for extensive large-scale fermentation and purification processes, thus offering a more streamlined approach to DNA-intercalating natural product discovery. By applying SMSA to 17 microorganisms, we identified two DNA intercalator-producing strains: *Streptomyces tanashiensis* and *Talaromyces funiculosus*. Subsequently, three DNA intercalators, namely medermycin, kalafungin, and ligustrone B, were isolated and characterized. Among them, medermycin and kalafungin showed significant inhibitory effects against HCT-116 cancer cells, with *IC*_*50*_ values of 52 ± 6 nM and 70 ± 7 nM, respectively.

## Introduction

DNA has long been a target for anticancer small-molecule medicines.^[1-2]^ Chemotherapeutic drugs that act as double-strand DNA (dsDNA)-binding ligands, including alkylating agents, intercalators and groove binders, remain a crucial pillar of many cancer treatment regimens.^[1]^ Approximately 17% (55 out of 321) of the therapeutic anticancer drugs approved between 1949 and 2019 are DNA interacting agents,^[3]^ with around half of these agents derived from natural products.^[3-4]^ Many notable anticancer natural products (antibiotics) are DNA intercalators, including actinomycin D, doxorubicin, idarubicin, and camptothecin.^[5]^ These intercalators bind DNA by inserting aromatic moieties between adjacent DNA base pairs, and subsequently disrupt fundamental processes like DNA replication and RNA transcription.^[6-7]^ Understanding of genome structure and function of genes has also created new opportunities for DNA intercalators targeting cellular DNA or the associated proteins, including topoisomerase inhibitors,^[8]^ and drug targeting non-canonical DNA structures like G-quadruplexes.^[9-10]^

Given the escalating issue of drug resistance, there is a pressing need for novel DNA interacting agents. Additionally, considering the diverse clinical efficacy observed among drugs from various chemical families targeting the same molecule, novel DNA-binding agents hold promise for combating different types of tumors. Microorganisms, which produce diverse secondary metabolites to gain a competitive advantage, represent a rich source of bioactive DNA-binding agents.^[11-12]^ However, the ability to produce DNA-interacting compounds is not universal among organisms, instead, such compounds are restricted to specific species under particular environmental conditions.^[12]^ This highlights the need for a more accessible and rapid screening method to identify microorganisms producing DNA intercalating compounds with chemical diversity.

The conventional approach to identifying microorganisms capable of producing bioactive natural products primarily relies on their biological activity in crude extracts or purified compounds.^[13]^ However, this approach may overlook potentially valuable medicinal compounds with low yields. Moreover, predicting the targets of these compounds before chemical purification and structural elucidation remains challenging. For the detection of DNA-binding compounds, the most commonly used and sensitive detection methods currently include surface plasmon resonance, ^[14]^ affinity mass spectrometry,^[15]^ fluorescence-based assays, ultraviolet spectroscopy (UV) ^[16]^ or circular dichroism (CD) methods. Although each of these techniques has specific advantages and limitations, they generally require purified reagents, especially for the analysis of weak non-covalent DNA-ligands interactions (binding affinity *K*a ∼10^5^ to 10^7^ M^-1^). However, the separation and purification process of small molecules necessitate a substantial culture volume (> 10 L of liquid culture or 1 kg of solid culture) to isolate more than a microgram of reagents.^[17]^

Direct manipulation of DNA at single-molecule level has enabled the measurement of its mechanical properties ^[18-19]^ and expanded our understanding of its interactions with proteins or small ligands.^[20-23]^ The worm-like chain model is commonly used to characterize DNA polymer using two parameters: the contour length (*L*0 = 0.34 nm) and the persistence length (A = 50 nm) to describe the force-extension curves of dsDNA.^[19]^ Previous studies have shown that each intercalator inserted into dsDNA leads to an increase in the contour length L0 of DNA base pairs by 0.2-0.3 nm.^[24]^ Moreover, the binding of intercalators and major groove binders can significantly alter the behavior of the DNA overstretching transition occurring at 65 pN.^[25]^ While the majority of previous single-molecule assays have been conducted with purified small molecules under defined conditions,^[21]^ the application of single-molecule manipulations to the field of natural product chemistry for the direct analysis of complex mixtures remains unexplored. It is not yet clear whether the SMSA is applicable in the presence of complex sample systems such as microbial cultures containing a wide variety of metabolites or proteins.

Here, we introduce a pioneering application of SMSA directly in microbial cultures or extracts to screen microorganisms that produce DNA intercalating agents (Fig. 1A). We demonstrated that a notable elongation of dsDNA (>15% of the extension of pure dsDNA) under forces (>10 pN) serves as a distinctive feature of DNA intercalators. Importantly, this elongation remains unaffected by the presence of primary metabolites involved in the basic growth of microorganisms, such as carbohydrates, lipids, and amino acids. Moreover, the application of external force on dsDNA can enhance the binding affinity and number of intercalations, thereby increasing the sensitivity of detection. The high sensitivity and specificity of the method enable the identification of trace amounts of DNA intercalative agents from just 5 μL of bacterial culture, eliminating the need for purification steps. It is noteworthy that the integration of magnetic tweezers with a microfluid flow-chamber allowed for buffer exchange, rendering it well-suited for measuring multiple samples in the same chamber.

**Figure 1.**
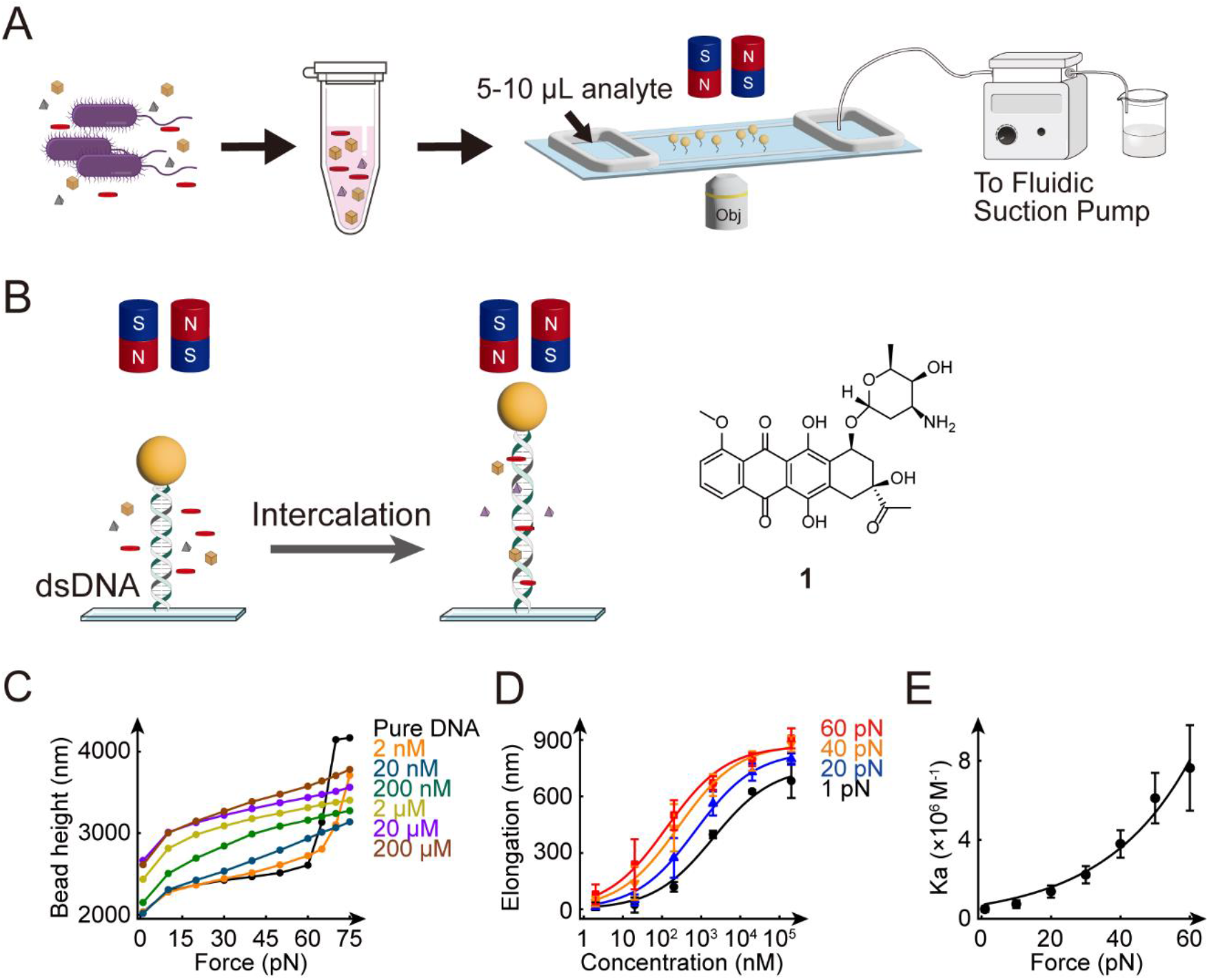
Experimental design. (A) Single-molecule stretching assay for microbial culture using magnetic tweezers. (B) A 6618 bp dsDNA tethered between a paramagnetic bead and a coverslip was used as a sensor to detect the presence of DNA intercalating agents. Chemical structure of daunorubicin (compound **1**). (C) Typical bead height-force curves measured in the presence of various daunorubicin concentration. The increase of bead height indicates the elongation of dsDNA due to intercalation by daunorubicin. (D) Titration curves of dsDNA elongation measured at different forces. Data points were expressed as the means ± standard deviations. Lines were fitted to the data by Hill function. (E) Force-dependent binding constants were approximated to the Arrhenius-Bell mode.

## Results and Discussion

### External forces increase the sensitivity and specificity for detecting DNA intercalating agents

For proof of principle, we employed a single-molecule magnetic tweezer to analyze a purified DNA intercalator, daunorubicin that is a widely used anthracycline antibiotic to treat various types of cancer.^[26]^ The experimental design using magnetic tweezers was shown in Fig. 1B, where a 6618 bp dsDNA was tethered between a paramagnetic bead and a cover glass slide, serving as a sensor to detect the DNA interacting agents.^[9, 21]^ By monitoring the extension of dsDNA using magnetic tweezers and applying forces ranging from 1 pN to 75 pN, the extension changes induced by DNA binding small molecule can be measured. The elongation of dsDNA measured in the presence of different concentration of daunorubicin was shown in Fig. 1C. Dissociation constants (*K*d) for daunorubicin at various forces were fitted using the Hill equation (Fig. 1D), 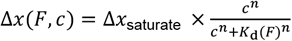, where Δ*x*(*F, c*) is the elongation of dsDNA at ligand concentration c and force *F*, Δ*x*_saturate_ is the saturate dsDNA elongation. *K*_d_ (F) is the force-dependent dissociation constant, and n is the Hill coefficient.

We observed a significant enhancement in the sensitivity for detecting DNA intercalators in response to applied force, which could be attributed to several factors. Firstly, DNA is resistant to being straightened in solution due to thermal fluctuations,^[19]^ which causes challenges in detecting intercalator-induced elongations. According to the worm-like chain model, a force exceeding *k*_B_*T*/*A* (> 0.1 pN) is needed to extend the dsDNA, where *k*_B_ is Boltzmann constants and *T* is the absolute temperature. The high spatial resolution of our magnetic tweezers (∼2 nm at forces >5 pN) allowed the detection of few molecules binding (0.34 nm/intercalation), thereby achieving sensitive detection. Secondly, the ligand-induced elongation is sensitive under high forces as force can increase the number of intercalators bound to dsDNA.^[24]^ For instance, in the presence of 20 nM daunorubicin, insignificant elongation was observed at low forces (<10 pN), whereas substantial elongation (329 ± 26 nm) occurred at 60 pN, representing approximately to 14±1% (fractional elongation) of the extension of pure dsDNA (Fig.1D). Thirdly, external forces applied to the dsDNA significantly enhance the binding affinity of ligands to dsDNA at force *F*, as determined by zero-force binding affinity *K*_a_(0) using the Arrhenius-Bell model 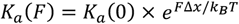,^[1]^ where Δ*x* is the transition distance. By the fitting of *K*_a_ measured at different forces to equation 1, we obtained zero-force *K*_a_ (0) = (0.71 ± 0.13) ×10^6^ M^-1^, and *Δx* = 0.21± 0.01 nm. The *K*_a_ increased ∼10-fold to (7.6 ± 2.1) × 10^6^ M^-1^ at 60 pN (Fig. 1E). The increase of DNA binding affinity under external force is a general property for dsDNA intercalators,^[23]^ thus this approach can be applied to wide range of unknown DNA intercalators, enabling the detection of low-abundance DNA intercalating agents from microorganisms.

In addition to lengthening dsDNA, the binding of ligands to dsDNA also altered the behavior of the DNA overstretching transition, providing a sensitive indicator for the presence of DNA interacting reagents.^[25, 27]^ At a critical force of ∼65 pN, pure dsDNA showed sudden 70% increase in extension to 0.58 nm per bp (Fig. 1B, black curve), indicating the transition of B-form DNA to S-DNA (stretched DNA). At 2 nM daunorubicin, the overstretching transition force increased, suggesting that 2 nM daunorubicin binds and stabilizes B-form DNA. The detection limit for strong DNA intercalator like daunorubicin is ∼20 nM, as the overstretching transition at ∼65 pN disappears in the presence of 20 nM daunorubicin. Given the small sample volume (5 μL) required for detection, SMSA can detect ligand-DNA interactions using only 0.052 ng of daunorubicin.

Importantly, the application of force also enhances the specificity for detecting DNA intercalators. This is because the lengthening of dsDNA at high force (>10 pN) is a specific characteristic of DNA intercalators.^[20]^ Solution conditions, primarily ionic strength, mainly influence the persistence length of dsDNA but do not significantly increase itscounter length.^[28]^ Therefore, changes in ionic strength do not lead to substantial elongation (less than 10% of the extension of pure dsDNA) at high force. Most DNA-binding proteins change the dsDNA extension by stiffening (increasing persistence length), bending or wrapping dsDNA, thereby reducing the extension at low force (<10 pN). However, they do not cause lengthening of dsDNA at high force.^[29]^ To our knowledge, only RecA and single-stranded DNA (ssDNA) form a filament that binds to dsDNA and can result in the lengthening of dsDNA.^[30]^ Therefore, we speculate that SMSA can directly detect DNA-intercalating agents from microbial culture.

### Validating the suitability of SMSA for detecting DNA intercalators in microbial culture using *S. coeruleorubidus*

To verify the suitability of SMSA for directly analyzing microbial culture, we investigated the liquid culture of the known daunorubicin-producing bacterium *S. coeruleorubidus*. Bacteria typically generate primary metabolites in the early stage and significant quantities of secondary metabolites during the late-exponential or stationary phases. By monitoring the absorbance at 480 nm, a characteristic wavelength for anthracyclines, we confirmed the presence of daunorubicin or its derivatives in the bacterial supernatants collected at various stages of culture growth (Fig. 2A). The dsDNA stretching curve with supernatants from early-phase bacterium culture (cultivated for 70 h) (Fig. 2B, red) and M2 medium (Fig. S1, red) overlapped with that of pure DNA (black), indicating no significant elongation. However, supernatants from late-exponential phase culture (cultivated for 178 h) induced significant dsDNA elongation (>450 nm) (Fig. 2C), suggesting the presence of DNA-intercalating agents. The significant elongation observed at forces ranging from 10 to 60 pN, accompanied by the disappearance of the B-to-S phase transition at 65 pN, is a characteristic of dsDNA intercalators. This result suggests that the dsDNA elongation at high forces (>10 pN) is highly specific to DNA intercalators, remaining unaffected by primary metabolites involved in the basic bacteria growth, such as carbohydrates, lipids, and amino acids. To broaden the scope of the applicability to various type of microorganism cultures, we also investigated single bacterium colonies from the agar plates cultured with *S. coeruleorubidus* (Fig. 2D). Each colony, along with the agar (weighing 135 ± 11 mg) was immersed in 1 mL of water and subjected to extraction using an ultrasonic bath sonicator for 30 minutes. Subsequently, 5 μL of the resulting supernatants were introduced into the flow chamber for SMSA. The bead height-force curve measured at force ranging from 10 pN to 60 pN exhibited significant elongation (ranging from 215 ± 77 nm to 499 ± 27 nm), indicating the presence of DNA intercalating agents in the bacterial colonies.

**Figure 2.**
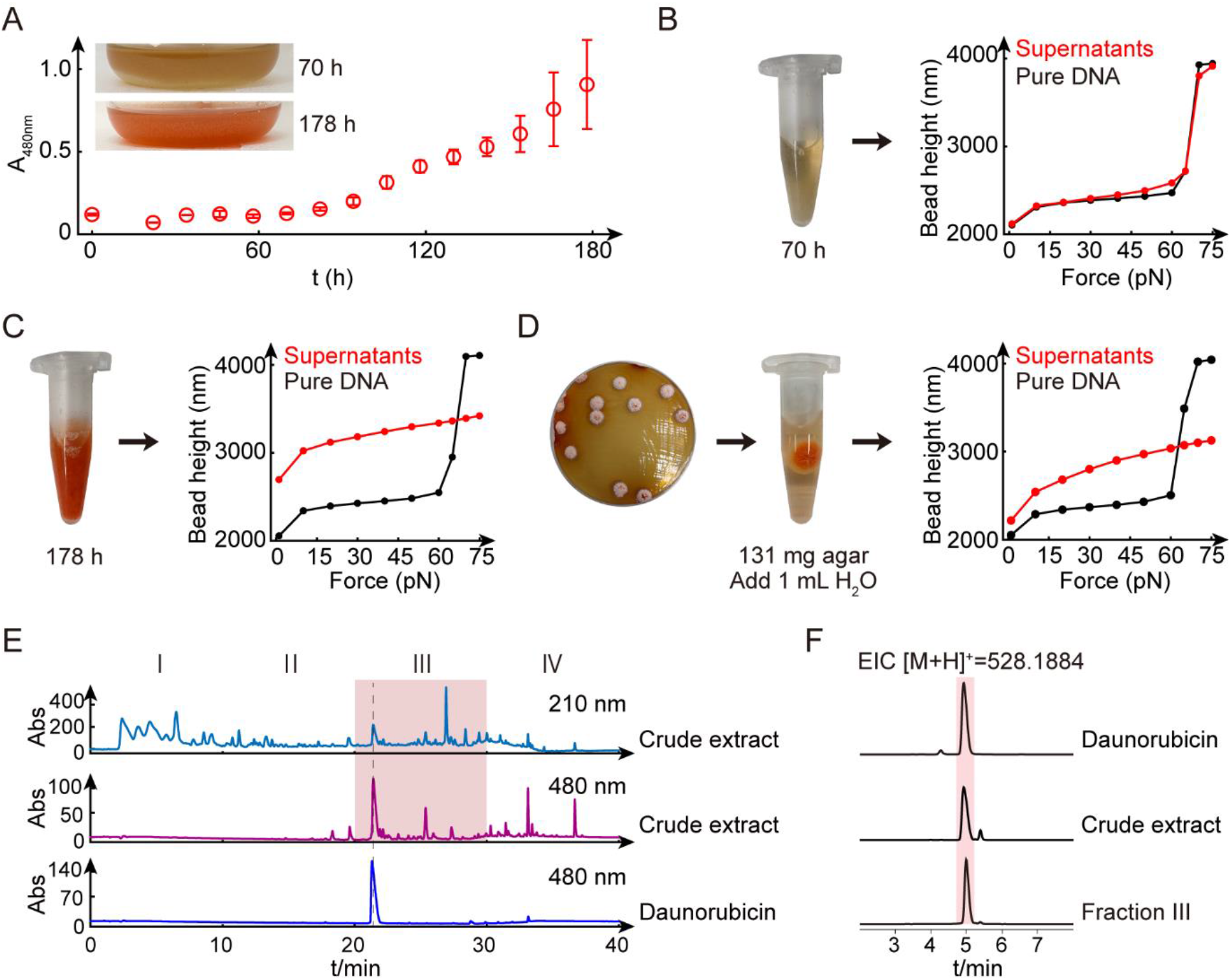
Validation of SMSA for analyzing culture of *Streptomyces coeruleorubidus*. (A) Production of daunorubicin and its derivatives was quantified based on the absorption at 480 nm. The data represent the mean values ± standard deviations obtained from three independent culture supernatants collected at 12-hour intervals. Additionally, photographs of culture flasks captured at 70 h and 178 h were included for reference. (B-C) SMSA results for the addition of 5 μL of bacterial supernatants from cultures cultivated for 70 h (B) and 178 h (C) were shown. The curves were color-coded: Black, pure DNA; Red, supernatants. (D) SMSA for bacterial colony. A single colony of *S. coeruleorubidus* grown on an M2 agar plate was selected and extracted with 1 mL of ddH_2_O, then 5 μL of the supernatants were subjected to the assay. (E) HPLC fragmentations of the crude extracts derived from *S. coeruleorubidus*. UV spectra of crude extracts at 210 nm (upper panel), 480 nm (middle panel), and daunorubicin standard (lower panel). (F) Extracted ion chromatogram (EIC) of (528.1884, [M+H]^+^) from daunorubicin standard, crude extracts and Fraction III.

Subsequently, high performance liquid chromatography (HPLC) was employed to fractionate the crude extracts (1 mg) of *S. coeruleorubidus* culture into four fractions at 10-minute intervals (Fig. 2E), enabling the separation of polar and nonpolar compounds. The resulting four fractions were subjected to the SMSA, with fraction III displayed the highest average elongation compared to the other fractions (Fig. S2). This suggests the presence of DNA binding reagents in fraction III, consistent with the retention time of the daunorubicin standard. Liquid chromatography-high resolution mass spectrometry (LC-HRMS) analysis further confirmed the presence of the molecular ion peak at *m/z* 528.1884 [M+H]^+^ for daunorubicin in both the crude extract and fraction III (Fig. 2F). Additionally, we observed the presence of DNA intercalating compounds in fraction II as well. This likely stems from daunorubicin derivatives produced by *S. coeruleorubidus*, as the biosynthetic pathway for daunorubicin also produces a variety of anthracycline derivatives.^[4]^ Taken together, our findings demonstrate that the SMSA can facilitate the identification of DNA intercalators from trace amounts of bacterial culture during chromatography purification without the need for isolating individual compounds. This is particularly valuable for unknown DNA-interacting agents lacking specific absorbance and molecular weight information.

### Identification of microorganisms producing DNA interacting agents using SMSA

It is noteworthy that the DNA interacting agents are not universally present in secondary metabolites of all organisms. To assess the prevalence of microorganisms capable of producing DNA intercalators across a diverse range, we employed the SMSA to analyze bacterial cultures from 17 distinct species, including representatives of Gram-positive bacteria, Gram-negative bacteria and fungi (Fig. 3A). The strains were classified based on the sequences of the Internal Transcribed Spacer (ITS) in fungi and 16S rRNA in bacteria (Table S1). Following cultivation of the strains, the supernatants from bacterial cultures and the ethyl acetate (EtOAc) crude extracts from fungi (1 mg dissolved in 0.1 mL DMSO or MeOH) were individually subjected to the same SMSA. The results revealed that the supernatants of *S. tanashiensis* DSM 731 exhibited the significant lengthening of dsDNA at forces > 30 pN, suggesting the presence of dsDNA intercalators in the culture (Fig. 3B). In contrast, all the analyzed Gram-negative bacterial cultures showed less than 10% fractional elongation at 60 pN (Fig. S3). Additionally, the extension of dsDNA measured in the presence of crude extracts produced by fungus *Talaromyces funiculosus* R5SSF2 contains DNA intercalators, evidenced by a significant elongation (943 ± 101 nm, fractional elongation = 42 ± 4%) of DNA at 60 pN (Fig. 3C), suggest that this species may also hold the potential for producing DNA intercalators.

**Figure 3.**
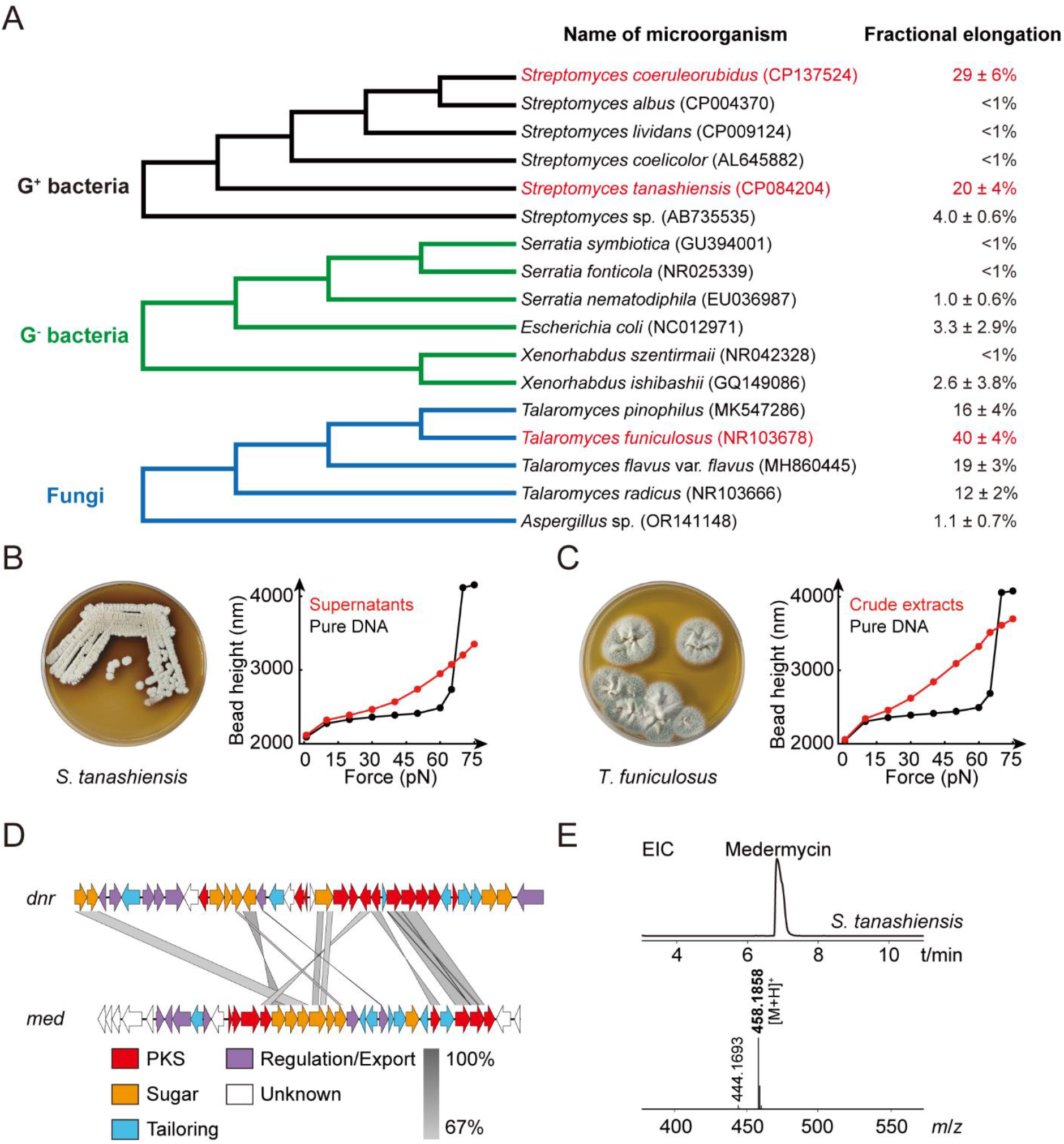
Screening of DNA intercalating agent producing microorganisms. (A) Phylogenetic analysis of the microorganisms used in this study. The culture supernatants of bacteria and the fungal crude extracts (0.1 mg/μL) were used for SMSA. Fractional elongation was estimated based on the elongation measured at 60 pN. Error represents standard deviations calculated from three independent measurements. The strains highlighted in red were used for subsequent analysis. (B-C) SMSA for the supernatants of *S. tanashiensis* DSM 731 (B) and the crude extract of *T. funiculosus* R5SSF2 (C). (D) The nucleotide comparison between the daunorubicin gene cluster (*dnr*) from *S. coeruleorubidus* and medermycin gene cluster (*med*) from *S. tanashiensis* generated using Easyfig 2.2.5. Grey shading indicates nucleotide similarity between two gene clusters (67 to 100%). Functioned regions of gene clusters were indicated: polyketide synthase (PKS): red, deoxysugar (Sugar): orange, post-PKS tailoring (Tailoring): blue, Regulation/Export: purple. (E) Extracted ion chromatogram (EIC) of medermycin (458.1858, [M+H]^+^) from the crude extract of *S. tanashiensis* DSM 731.

Natural products are synthesized by specific gene clusters within the genomes of their respective organisms. To preliminarily identify the types of DNA-interacting agents produced by *S. tanashiensis* DSM 731, we analyzed its complete genome sequence (CP084204) using antiSMASH (The antibiotics and Secondary Metabolites Analysis Shell).^[31]^ The analysis revealed the presence of the medermycin biosynthetic gene cluster (*med*), responsible for producing medermycin, an anthracycline polyketide. ^[32]^ Daunorubicin, another typical anthracycline polyketide, is encoded by a type II polyketide synthase (PKS) gene cluster known as *dnr*. ^[33]^ Comparison of the *dnr and med* clusters showed a similarity of at least 67%. These gene clusters can be categorized into four parts: PKS related genes for anthracycline core (PKS), deoxysugar genes (Sugar), the tailoring genes for methylation and oxidation (Tailoring), and regulatory genes (Regulation/Export) (Fig. 3D). Based on these findings, we hypothesized that *S. tanashiensis* DSM 731 can produce medermycin in bacteria supernatants, leading to the elongation in SMSA. To verify this hypothesis, we analyzed crude extracts of *S. tanashiensis* DSM 731 using LC-HRMS. The extracted ion chromatogram (EIC) showed a single prominent peak with an *m/z* value of 458.1858 ([M+H]^+^) (Fig. 3E), suggesting that *S. tanashiensis* DSM 731 produces medermycin. However, we were unable to perform biosynthetic gene cluster analysis (genome mining) for the fungus *T. funiculosus* due to the unavailability of its genome.

### Purification and characterization of DNA intercalators from the Gram-positive bacterium and the fungus

To further characterize the DNA intercalating compounds from the cultures of the Gram-positive bacterium *S. tanashiensis* DSM 731 and the fungus *T. funiculosus*, we conducted large-scale fermentation for these two strains. Guided by SMSA, we purified the target DNA intercalators, resulting in the isolation of compounds **2**-**4**, respectively. Subsequent analysis using ^1^H and ^13^C NMR spectroscopic data, as well as high-resolution electrospray ionisation mass spectrometry (HRESIMS) data revealed that compounds **2**-**3** isolated from *S. tanashiensis* DSM731 were identified as medermycin and kalafungin,^[34]^ originally discovered from *Streptomyces* sp. K73 in 1976. Compounds **2**-**3** belong to the family of pyranonaphthoquinone antibiotics, known for their antibacterial and anticancer bioactivity. Compound **4**, isolated from fungi *T. funiculosus* was structurally characterized as the known compound ligustrone B.

Using SMSA, we further individually analyzed the interactions of purified compounds **2**-**4** (Fig. 4A). The force-extension curves of dsDNA at serial concentrations of compounds **2**-**4** showed the concentration-dependent elongation from 1 to 60 pN (Fig. 4B). Notably, the overstretching transitions were significantly influenced by compounds **2**-**4** at concentrations beyond 10 μM. The elongation−concentration curves of compounds **2**-**4** were fitted with the Arrhenius-Bell model (Fig. S4). This allowed us to obtained the zero-force binding constants *K*a of compounds **2**-**4** when intercalating into dsDNA: (3.5 ± 0.3) × 10^3^ M^-1^ (**2**), (1.8 ± 0.2) × 10^3^ M^-1^ (**3**) and (10.5 ± 1.7) × 10^3^ M^-1^ (**4**), respectively. It is noteworthy that stretching curves under low force reveals distinct intercalating features among these compounds. In comparison to compound **3**, the elongation of dsDNA induced by compound **2** showed a more pronounced increase when measured at forces below 15 pN, which suggest higher number of ligands intercalation at low forces. This implies that the electrophilic side chain of compound **2** aids the molecule in inserting between the bases in dsDNA at low forces.

**Figure 4.**
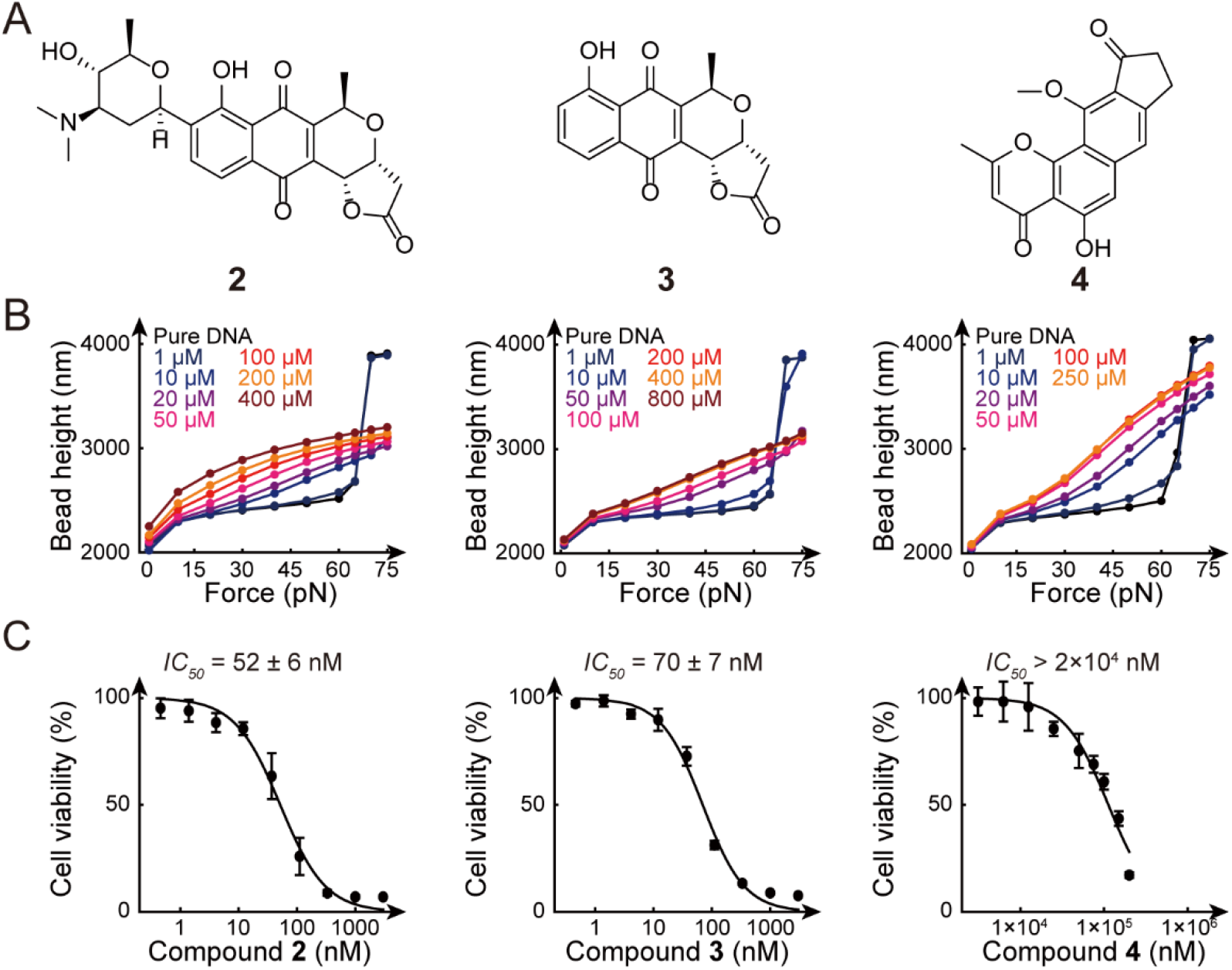
Characterization of purified DNA interacting agents. (A) The chemical structures of medermycin (**2**), kalafungin (**3**), ligustrone B (**4**). (B) Bead height-force curves measured in the presence of various concentrations of compounds **2**-**4**. (C) *IC*_*50*_ value of compounds **2**-**4** measured in HCT-116 cells. The y-axis represents relative cell viability compared to non-treated cells (control). The error bars represent the standard deviations from three independent replicates.

We also assessed the cytotoxic potential of compounds **2-4** against human colon cancer HCT-116 cells using MTT assay to measure cell proliferation. Medermycin and kalafungin exhibited remarkable antiproliferative effects against HCT-116 cells, with half-maximal inhibitory concentration (*IC*_*50*_) of 52 ± 6 nM and 70 ± 7 nM, respectively. In contrast, compound **4** showed only very weak activity against HCT-116 cells with *IC*_*50*_ greater than 2×10^4^ nM (Fig. 4C). The significant activity of compound **2** is consistent with previous reports, emphasizing its therapeutic potential in cancer treatment.^[35]^ The low activity of compound **4** is likely attributed to its low solubility, which could impede its cellular uptake and target engagement.

## Conclusion

In this study, we present a highly sensitive and specific single-molecule assay that enables the direct detection of DNA intercalating ligands from minute amounts of microorganism cultures. This mechanical assay is highly specific for DNA intercalators and is unaffected by the presence of various metabolites, eliminating the need for complex sample separation or purification. Our assay required only nanogram of DNA interacting reagents, less than 1% of the amount needed for any other methods, facilitating rapid screening of microorganisms producing DNA intercalating reagents with low abundance.

One of the major challenges in conventional natural product discovery is the laborious workflow involved in fermenting and isolating natural products from microbes, often resulting in the rediscovery of known compounds. Advances in high-throughput sequencing and mass spectroscopy technologies have made massive data on microbial genomes, transcriptomes and proteomes readily accessible. While bioinformatic tools excel at identifying conserved biosynthetic gene clusters responsible for the production of compounds, such as polyketides, non-ribosomal peptides, terpenes, etc.,^[31]^ they fall short in predicting the actual DNA binding properties of secondary metabolites. In this context, integrating SMSA with genome mining strategies and mass spectroscopy analysis streamlines the identification process for new chemical scaffolds with DNA-binding properties.

In our study, the isolated compounds medermycin and kalafungin showed modest DNA intercalating activity but displayed significant inhibition of cancer cell proliferation at sub-micromolar concentrations (Fig. 4). This implies that the force-enhanced binding affinity facilitated by SMSA enables the identification of weak DNA intercalators from microorganism cultures, while still exhibiting potent bioactivity. This may be attributed to the fact that many DNA-interacting drugs also interfere with DNA-associated proteins such as topoisomerase,^[8]^ RNA polymerase and telomerase. For instance, etoposide, a clinically used camptothecin derivative acting as a topoisomerase I poison, has been demonstrated to intercalate into bare DNA.^[36]^ Unlike typical DNA intercalators like ethidium bromide, topotecan acts as a weak dsDNA intercalator,^[36]^ but it is found at the interfaces between DNA and TOP1, forming a ternary complex.^[8]^ The discovery of novel DNA intercalators may contribute to the identification of novel non-camptothecin TOP1 inhibitors. Additionally, RNA intercalators also have a wide range of applications, exemplified by the first approved small molecule treatment for Spinal Muscular Atrophy (SMA), which showed intercalative binding to RNA hairpins.^[37]^ By substituting the dsDNA tether with dsRNA or RNA/DNA hybrid tethers, the current approach can also be adapted to detect RNA-interacting small molecules from microorganisms. ^[38]^

Advanced single-molecule techniques such as magnetic tweezers, optical tweezers, and nanopore sensors hold substantial promise for natural product discovery.^[39-40]^ These methodologies allow for precise and targeted measurements of various interactions, including those between ligands and DNA or protein.^[41]^ While it is regrettable that we did not discover any DNA-interacting agents with new chemical scaffolds due to the limited numbers of microorganisms screened, our findings still underscore the effectiveness of SMSA in specifically detecting DNA-interacting agents. By refining microfluidic systems and implementing automation, the throughput of SMSA can be significantly improved, thus expanding its potential applications in drug discovery endeavors.

## Supporting information

Supplemental experimental procedures, supplemental Table 1, supplemental Figure S1-S22

## Supporting Information

Supporting Information includes materials, methods, experimental procedures, other biological results, structural characterization of compounds, ^1^H, ^13^C NMR spectra as well as UV and HRESIMS spectra of compounds. The data underlying this study are available in the published article and its Supporting Information.

## Acknowledgements

This work was financially supported by the National Natural Science Foundation of China (No. 32171225, 32371496, 31972852 and 81903527) and the key project at central government level: The ability establishment of sustainable use for valuable Chinese medicine resources (No. 2060302). The authors appreciate the help from technicians at the Analytical and Testing Centre of the Huazhong University of Science and Technology with the NMR measurements.

## Entry for the Table of Contents

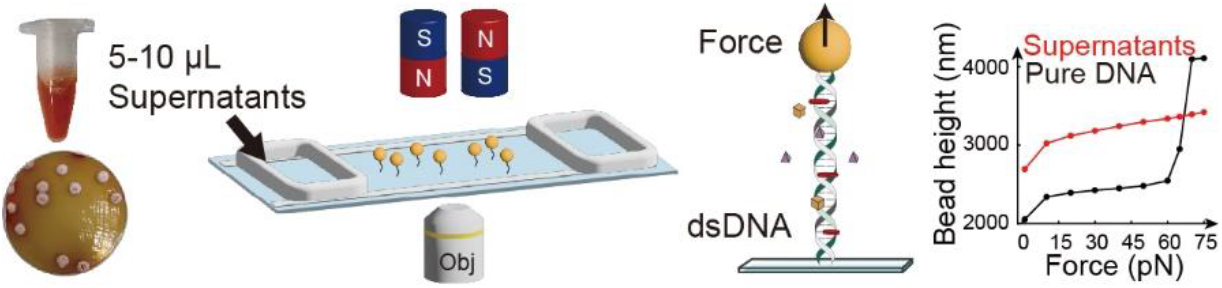

We present a novel application of single-molecule stretching assay using magnetic tweezers for direct detection of DNA-intercalating agents from complex microbial cultures. The alterations triggered by intercalators in the double-stranded DNA counter length and overstretching transition generate a distinct signature for DNA binding, facilitating the easy detection of DNA intercalators in trace amounts of microbial cultures.

