## Supplemental experimental procedures, supplemental Table 1, supplemental Figure S1-S22 for "Force-Enhanced Sensitive Detection of New DNA-Interactive Agents from Microorganisms at the Single-Molecule Level"

Angewandte Chemie

Eine Zeitschrift der Gesellschaft Deutscher Chemiker

[b] Prof. X. Cai
State Key Laboratory of Dao-di Herbs, Beijing 100700, China

[c] Prof. H. Liu
State Key Laboratory of Mycology, Institute of Microbiology, Chinese Academy of Sciences, Beijing 100101, China

[d] Prof. A. Li
Helmholtz International Lab for Anti-Infectives, Shandong University-Helmholtz Institute of Biotechnology, State Key Laboratory of Microbial Technology, Shandong University, Qingdao 266237, China

[†] These authors contributed equally to this work.

**Figure S15.** UV spectrum of compound **4**. 15
**Figure S16.** HRESIMS spectrum (positive ionization) of compound **4**. 15
**Figure S17.** $^1$H NMR (400 MHz, $CDCl_3$) spectrum of compound **4**. 16
**Figure S18.** $^{13}$C NMR (100 MHz, $CDCl_3$) spectrum of compound **4**. 16
**Figure S19.** DEPT-135° spectrum of compound **4**. 17
**Figure S20.** HSQC spectrum for compound **4**. 17
**Figure S21.** HMBC spectrum for compound **4**. 18
**Figure S22.** $^1$H-$^1$H COSY spectrum for compound **4**. 18
**References** 19

### Experimental procedures

#### 1. Reagents

Daunorubicin hydrochloride was purchased from Shanghai yuanye Bio-Technology Co., Ltd (China). The chemical solvents including petroleum ether, ethyl acetate, dichloromethane and methanol used for extraction and column chromatography were obtained from Hubei Shenshi Chemical Technology Co., Ltd (China). The HPLC-grade solvents (methanol and acetonitrile) used for LC and LC/MS analysis were procured from Fisher Chemical/Merck (USA/GER). Trifluoroacetic acid (TFA, HPLC-grade) was acquired from Energy Chemical (China). Deuterated solvents, namely, methanol-$d_4$ and chloroform-*d* were sourced from Cambridge Isotope Laboratories, Inc (UK).

#### 2. Strains and culture conditions

The strain of *S. coeruleorubidus* CICC11043 was acquired from China Center of Industrial Culture Collection. *S. tanashiensis* DSM 731, *Serratia symbiotica* DSM 23270, *S. fonticola* DSM 4576, *S. nematodiphila* DSM 21420, *Xenorhabdus szentirmaii* DSM 16338 and *X. ishibashii* DSM 22670 were obtained from Leibniz Institute DSMZ-German Collection of Microorganisms and Cell Cultures GmbH. *Streptomyces* sp. Gö66 was generously provided by Prof. Dr. Axel Zeeck from the University of Göttingen, Germany. *S. coelicolor* A3(2), *S. lividans* K4-114, *S. albus* J1074 and *Escherichia coli* BL21(DE3) are general hosts maintained in our lab. *Talaromyces pinophilus* R5SSF1, *T. funiculosus* R5SSF2, *T. flavus* var. *flavus* R5HwF7 and *T. radicus* BLYF1 were isolated from *Lycium barbarum* at 105°97'E, 38°28'N, Hui Autonomous Region, Ningxia, China. *Aspergillus* sp. CarHF15 was obtained from *Curcuma aromatica* Salisb at 116°25'E，39°47'N, Nanning, Guangxi, China.

All fungal strains were individually inoculated onto potato dextrose agar (PDA) plates and incubated at 28°C for 7 days. Subsequently, the fungus-growing PDA agar was cut into small pieces and transferred into sterilized rice media composed of 120 g of rice and 180 mL of double-distilled water (dd $H_2O$) at 28 °C for 20 days. For Gram-positive bacteria, cells were initially cultivated on GYM agar plates (1L: 4 g of d-glucose, 4 g of yeast extract, 10 g of malt extract, 2 g of $CaCO_3$, pH 7.2, and 20 g of agar) at 30 °C for 5 days for sporulation. Subsequently, the spore suspension of cells was transferred into M2 media (1L: 4 g of d-glucose, 4 g of yeast extract, 10 g of malt extract, pH 7.0, and 20 g $L^{-1}$ of agar for solid cultivation) at 30 °C, 200 rpm for 7 days, except for *S. tanashiensis* DSM 731 which was inoculated into R4 medium (1L: 10 g of d-glucose, 10 g of $MgCl_2·6H_2O$, 5.6 g of 2-[tris(hydroxymethyl)methylamino]-1-ethanesulfonic acid, 4 g of $CaCl_2·2H_2O$, 3 g of l-proline, 1 g of yeast extract, 0.2 g of $K_2SO_4$, 0.1 g of casamino acids, 2 mL of trace element solution, pH 7.2, and 20 g $L^{-1}$ of agar for solid cultivation). The trace element solution was prepared by dissolving 40 mg of $ZnCl_2$, 200 mg of $FeCl_3·6H_2O$, 10 mg of $CuCl_2·2H_2O$, 10 mg of $MnCl_2·4H_2O$, 10 mg of $Na_2B_4O_7·10H_2O$ and 10 mg of $(NH_4)_6Mo_7O_{24}·4H_2O$ were dissolved in 1 L of dd $H_2O$. Six Gram-negative bacteria were separately cultured on Luria-Bertani (LB) agar plates (1L: 10 g of tryptone, 5 g of yeast extract, 10 g of NaCl, pH 7.2, and 20 g of agar) and subsequently inoculated into trypticase soya broth (TSB) media for fermentation (1L: 30 g of TSB powder) at 30 °C, 200 rpm for 3 days.

#### 3. Single-molecule stretching assay

SMSA utilized single-molecule magnetic tweezers (BioPSI, Singapore). The measurements were conducted with a 6618 bp PCR product amplified from bacteriophage DNA (λDNA) (Thermo Fisher Scientific, USA) using a pair of primers labeled with biotin and thiol group at the respective 5' end (Sangon Biotech Co., Ltd, China). A flow-channel was prepared as previously described[1]. The 5'-thiol end of dsDNA was attached to the coverslip surface via a sulfosuccinimidyl 4-(*N*-maleimidomethyl) cyclohexane-1-carboxylate (sulfo-SMCC) crosslinker (Huateng Pharma Co., Ltd, China), while the 5'-biotin-end of dsDNA was attached to 2.8 µm-diameter streptavidin-coated paramagnetic beads Dynal M280 (Thermo Fisher Scientific, USA). The force was applied to the magnetic beads by a pair of permanent magnets positioned above sample. The bead-height was measured from the diffraction patterns of the beads using magnetic tweezers. The bead-height force curves were obtained through a force-jump procedure, and at each force, the magnet was held for 5 s to measure the bead-heights. All single-molecule experiments were conducted at room temperature of 20-23°C.

**4. Bacterial samples preparation for single-molecule stretching assay**

A 0.3 mL of bacterial liquid culture was centrifugated at 12000 rpm for 5 min and the supernatant was heated at 95°C for 5 min to inactivate DNase before SMSA. For agar cultures, the bacterial colony was picked from the agar plate and extracted with 1 mL of dd$H_2O$ and centrifugated at 12000 rpm for 5 min to collect the supernatant for SMSA. For crude extracts from fungi, 50 g rice culture was extracted with 50 mL of EtOAc and the solvent was evaporated under vacuum to obtain the crude extract. The crude extract from each microorganism was accurately weighed and dissolved in methanol to ensure consistent concentration (1 mg/ml). The dissolved crude extract was diluted 100-fold into 1 × PBS buffer (pH7.3) before SMSA. Finally, 5 µL of the analyte was added to the flow chamber.

**5. HPLC guided fractioning**

The crude extract of *S. coeruleorubidus* was dissolved in 500 µL of MeOH and subsequently analyzed using HPLC. The analysis was performed employing an Agilent Technologies 1260 Infinity system equipped with an Agilent Proshell 120 EC-$C_{18}$ column (2.7 µm, 150 × 3.0 mm). Liquid chromatography was conducted with acetonitrile containing 0.01% TFA (v/v) as solution A and milliQ $H_2O$ containing 0.01% TFA (v/v) as solution B. A linear gradient from 10% to 45% A was applied over 0 to 25 min, followed by a rapid gradient from 45 to 100% A from 25 to 30 min. The column was washed for 10 min with 100% A and then equilibrated with an isocratic flow at 10% A from 40 to 45 min. Chromatography of compounds was continuously monitored by measuring absorbance at 210 and 480 nm. To determine the retention time of daunorubicin, the organic extract materials were fractionated into 4 fractions at a flow rate of 0.6 mL/min, with each fraction collected for 10 min per tube. All tubes containing HPLC fractions were dried using a vacuum centrifugal concentrator (Beijing JM Technology, China). After drying, 100 µL of methanol was added to each fraction and then diluted 100 times with 1 × PBS solution. The prepared samples (100 µL) were added to the channel for SMSA.

**6. LC-MS analysis**

The filtered samples were analyzed by an Agilent & 1290 Infinity II / 6545 QTOF LC/MS using an Agilent Zorbax Eclipse Plus C18 RRHD column (1.8 µm, 50 × 2.1 mm) with a linear gradient of 5% to 100% solvent A (solvent A: 0.1% HCOOH in $CH_3CN$; solvent B: 0.1% HCOOH in $H_2O$) over 17 min at a flow rate of 0.4 mL/min.

**7. Phylogenetic and gene cluster analysis**

The sequences used in this study were obtained from the National Center for Biotechnology Information (NCBI, www.ncbi.nlm.nih.gov/) and analyzed using the BLAST algorithm. The phylogenetic tree was constructed via the maximum likelihood algorithm implemented in MEGA7.0, based on multiple sequence alignment by MUSCLE.

Bootstrap values were calculated after 1000 replications. The accession numbers of selected sequences were listed in Table S1. The nucleotide comparison of gene cluster generated using Easyfig 2.2.5[2].

**8. Purification and structural elucidation of DNA intercalators**

*S. tanashiensis* DSM 731 was grown in 5L of R4 media for 7 days at 30 °C with shaking at 200 rpm. The culture was then extracted with ethyl acetate four times at room temperature and the solvent was evaporated under vacuum to obtain 586.0 mg of crude extract. This extract was subjected to silica gel column chromatography (100-200 and 200-300 mesh, Qingdao Marine Chemical Inc., Qingdao, China) and eluted with a gradient of petroleum ether-ethyl acetate (v/v 10:1, 5:1, 1:1, 1:5, 1:10, 0:1) and $CH_2Cl_2$-MeOH (v/v 20:1, 10:1, 1:1, 0:1) to afford eight fractions (Fractions A-H). Fraction D (51.2 mg) was further purified through semipreparative HPLC (equipped with a Cosmosil 5$C_{18}$-MS-Ⅱ column, 5 µm, 250 × 10 mm) using 50% acetonitrile in $H_2O$ as the mobile phase, resulting in the isolation of compound **3** (7.6 mg, $t_R$ 34.280 min). Fraction G (129.8 mg) was separated by Sephadex LH-20 column chromatography (Cytiva, $CH_2Cl_2$:MeOH 1:1), followed by purification using semipreparative HPLC with isocratic elution (18% Acetonitrile in $H_2O$ ) to yield compound **2** (4.5 mg, $t_R$ 32.084 min) (Figure S5).

*T. funiculosus* R5SSF2 was grown on 480 g of rice media for 20 days at 28 °C. The fermented rice was then extracted with EtOAc three times at room temperature, and the solvent was evaporated under vacuum to obtain 10.2 g of crude extract. This extract was chromatographed on a silica gel column, eluted with a gradient of petroleum ether- EtOAc (v/v 10:1, 5:1, 1:1, 1:5, 1:10, 0:1) to give six fractions (Fractions A-F). Fraction C (1.7 g) was separated by Sephadex LH-20 column chromatography $CH_2Cl_2$:MeOH 1:1) to afford four fractions (C1-C4). Fraction C2 (0.6 g) was fractionated by silica gel column chromatography (eluted with a gradient of petroleum ether- EtOAc from 10:1 to 1:10, v/v) to yield five fractions (C2-1 - C2-5). Fraction C2-4 (51.2 mg) was further purified by semipreparative HPLC with 55% acetonitrile in $H_2O$ as the mobile phase to obtain compound **4** (7 mg, $t_R$ 29.578 min) (Figure S5).

All nuclear magnetic resonance (NMR) data were recorded on a Bruker AM-400 spectrometer with TMS as the internal standard. The structures of compounds **2**-**4** were elucidated by analysis of their spectroscopic data (NMR and HRESIMS, Figure S6-S22) and were corroborated with literature values[3, 4, 5].

Medermycin (compound **2**): red brown solid; HRESIMS *m/z* 458.1858 $[M+H]^+$ (calcd for $C_{24}H_{28}NO_8$, 458.1809); $^1H$ NMR (400 MHz, $CD_3OD$) $\delta$ 7.92 (d, $J$ = 7.8 Hz, 1H, H-7), 7.67 (d, $J$ = 7.8 Hz, 1H, H-6), 5.31 (d, $J$ = 2.9 Hz, 1H, H-4), 5.06 (q, $J$ = 6.9 Hz, 1H, H-1), 5.01 (dd, $J$ = 10.8, 2.1 Hz, 1H, H-1'), 4.78 (dd, $J$ = 5.2, 2.9 Hz, 1H, H-3), 3.63 (m, 1H, H-3'), 3.57 (m, 1H, H-5'), 3.48 (dd, $J$ = 10.2, 8.6 Hz, 1H, H-4'), 3.16 (dd, $J$ = 17.8, 5.2 Hz, 1H, H-11), 2.95 (s, 3H, 3'-NMe), 2.83 (s, 3H, 3'-NMe), 2.56 (m, 2H, H-2', 11), 1.69 (q, $J$ = 12.2, 1H, H-2'), 1.56 (d, $J$ = 6.9 Hz, 3H, H-13), 1.42 (d, $J$ = 6.0 Hz, 3H, H-6'); $^{13}C$ NMR (100 MHz, $CD_3OD$) $\delta$ 190.02 (C-10), 182.84 (C-5), 177.27 (C-12), 158.87 (C-9), 150.99 (C-10a), 137.84 (C-8), 136.75 (C-4a), 134.71 (C-7), 132.12 (C-5a), 120.02 (C-6), 115.85 (C-9a), 78.58 (C-5'), 72.33 (C-1'), 71.21 (C-4'), 70.83 (C-4), 68.53 (C-3'), 68.09 (C-3), 67.71 (C-1), 42.21 (3'-NMe), 37.65 (C-11), 37.47 (3'-NMe), 30.29 (C-2'), 18.51 (C-13), 18.19 (C-6').

Kalafungin (compound **3**): orange yellow solid; HRESIMS *m/z* 301.0708 $[M+H]^+$ (calcd for $C_{16}H_{13}O_6$, 301.0707); $^1H$ NMR (400 MHz, $CDCl_3$) $\delta$ 11.83 (s, 1H, 9-OH), 7.67 (m, 2H, H-6, 7), 7.29 (dd, $J$ = 7.3, 2.3 Hz, 1H, H-8), 5.25 (d, $J$ = 2.8 Hz, 1H, H-4), 5.08 (q, $J$ = 6.9 Hz, 1H, H-1), 4.69 (dd, $J$ = 5.2, 2.8 Hz, 1H, H-3), 2.97 (dd, $J$ = 17.7, 5.2 Hz, 1H, H-11), 2.69 (d, $J$ = 17.7 Hz, 1H, H-11), 1.56 (d, $J$ = 6.9 Hz, 3H, H-13); $^{13}C$ NMR (100 MHz, $CDCl_3$) $\delta$ 188.11 (C-10), 181.63 (C-5), 174.12 (C-12), 162.02 (C-9), 149.88 (C-10a), 137.33 (C-7), 135.25 (C-4a), 131.58 (C-5a), 125.00 (C-8), 119.87 (C-6), 114.93 (C-9a), 68.77 (C-4), 66.59 (C-3), 66.38 (C-1), 37.02 (C-11), 18.70 (C-13).

Ligustrone B (compound **4**): yellow needle crystal; HRESIMS *m/z* 311.0940 $[M+H]^+$ (calcd for $C_{18}H_{15}O_5$, 311.0914); $^1$H NMR (400 MHz, $CDCl_3$) $\delta$ 12.94 (s, 1H, 6-OH), 7.41 (s, 1H, H-9), 6.95 (s, 1H, H-7), 6.38 (s, 1H, H-3), 4.16 (s, 3H, 15-OMe), 3.23 (m, 2H, H-11), 2.78 (m, 2H, H-12), 2.57 (s, 3H, 2-Me); $^{13}$C NMR (100 MHz, $CDCl_3$) $\delta$ 203.47 (C-13), 182.94 (C-4), 167.53 (C-2), 158.49 (C-6), 158.42 (C-15), 157.43 (C-17), 154.23 (C-10), 143.73 (C-8), 124.23 (C-14), 118.90 (C-9), 111.73 (C-16), 111.32 (C-3), 110.95 (C-5), 106.41 (C-7), 63.01 (15-OMe), 37.47 (C-12), 25.36 (C-11), 20.71 (2-Me).

**9. Cytotoxicity Assay**

The HCT-116 human colon cancer cells were a generous gift from Prof. Shunichi Takeda. The HCT-116 cells were initially cultured in high-glucose DMEM (Wuhan Servicebio Technology Co., Ltd) supplemented with 10% of fetal bovine serum (FBS, Pan-Biotech) and 100 U/mL penicillin/streptomycin (Beyotime Biotechnology, China). Cells were seeded at a density of 3000 cells/well in a 96-well plate and incubated for 24 h. Subsequently, cells were treated with the test compounds at different concentrations. After 48 h incubation at 37 °C, MTT (3-(4,5-dimethylthiazol-2-yl)-2,5-diphenyltetrazolium bromide, BioFroxx, Germany) in 1× PBS was added to each well at a final concentration of 0.25 mg/mL. The plate was further incubated for 2 h, followed by replacement of the medium with 100 µL of DMSO to solubilize the formazan products. The absorbance of the wells at 570 nm was measured using a Synergy H1 microplate reader (BioTek Instruments, Inc., USA). The $IC_{50}$ value was determined by fitting the relative cell viability to $y = \frac{100}{1+10^{M*(LogIC50-Logc)}}$.

### Supplementary Tables

**Table S1.** 16S rRNA and ITS sequences of microorganisms used in this study.

| No | Name of microorganism | Type of sequences | Length, bp | Accession |
|---|---|---|---|---|
| 1 | *S. coeruleorubidus* CICC 11043 | 16S rRNA | 1528 | CP137524 |
| 2 | *S. tanashiensis* DSM 731 | 16S rRNA | 1526 | CP084204 |
| 3 | *Streptomyces* sp. Gö66 | 16S rRNA | 1517 | AB735535 |
| 4 | *S. coelicolor* A3(2) | 16S rRNA | 1531 | AL645882 |
| 5 | *S. lividans* K4-114 | 16S rRNA | 1515 | CP009124 |
| 6 | *S. albus* J1074 | 16S rRNA | 1515 | CP004370 |
| 7 | *S. symbiotica* DSM 23270 | 16S rRNA | 1478 | GU394001 |
| 8 | *S. fonticola* DSM 4576 | 16S rRNA | 1490 | NR025339 |
| 9 | *S. nematodiphila* DSM 21420 | 16S rRNA | 1500 | EU036987 |
| 10 | *X. szentirmaii* DSM 16338 | 16S rRNA | 1524 | NR042328 |
| 11 | *X. ishibashii* DSM 22670 | 16S rRNA | 1480 | GQ149086 |
| 12 | *E. coli* BL21(DE3) | 16S rRNA | 1542 | NC012971 |
| 13 | *T. pinophilus* R5SSF1 | ITS | 620 | MK547286 |
| 14 | *T. funiculosus* R5SSF2 | ITS | 612 | NR103678 |
| 15 | *T. flavus* var. *flavus* R5HwF7 | ITS | 577 | MH860445 |
| 16 | *T. radicus* BLYF1 | ITS | 636 | NR103666 |
| 17 | *Aspergillus* sp. CarHF15 | ITS | 578 | OR141148 |

### Supplementary Figures

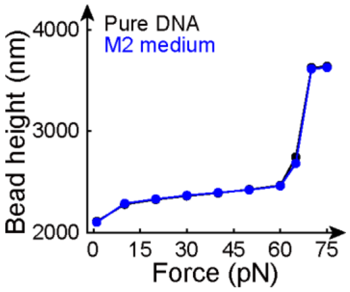

**Figure S1.** Single-molecule assay of M2 medium. Black, pure DNA; Blue, after addition of 5 µL M2 medium.

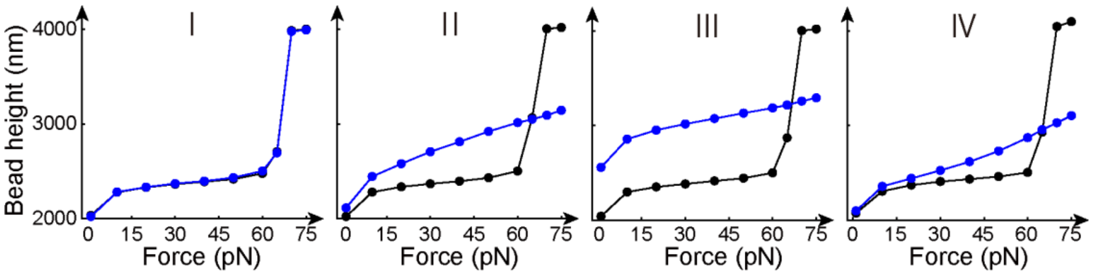

**Figure S2.** Single-molecule assay of fractions I to IV from HPLC separation of crude extracts of *S. coeruleorubidus.* HPLC fractions were dried and then dissolved in 100 µL methanol. Black, pure DNA; Blue, after addition of 100 µL prepared sample containing 1 µL methanol solution and 99 µL PBS buffer.

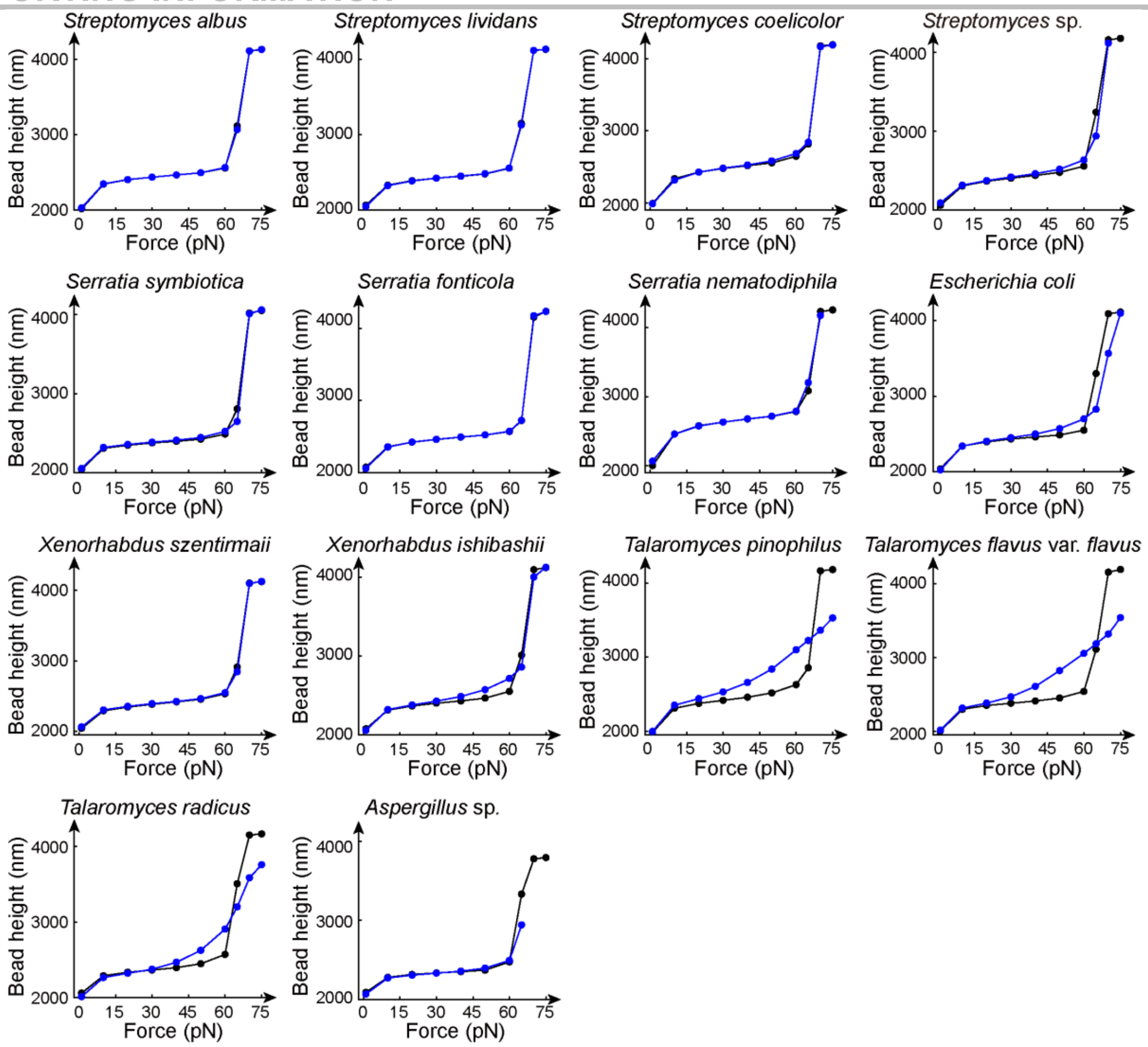

**Figure S3.** Single-molecule screening of microbial strains. Black, pure DNA; Blue, after addition of prepared samples. For Gram-positive bacteria (*Streptomyces* strains) and Gram-negative bacteria (*Serratia*, *Escherichia* and *Xenorhabdus* strains), 5 μL culture supernatants were subjected to the channel for the single-molecule stretching assay. For fungi (*Talaromyces* and *Aspergillus* strains), 100 μL crude extracts (0.1 mg/mL) were added to the channel for the single-molecule stretching assay.

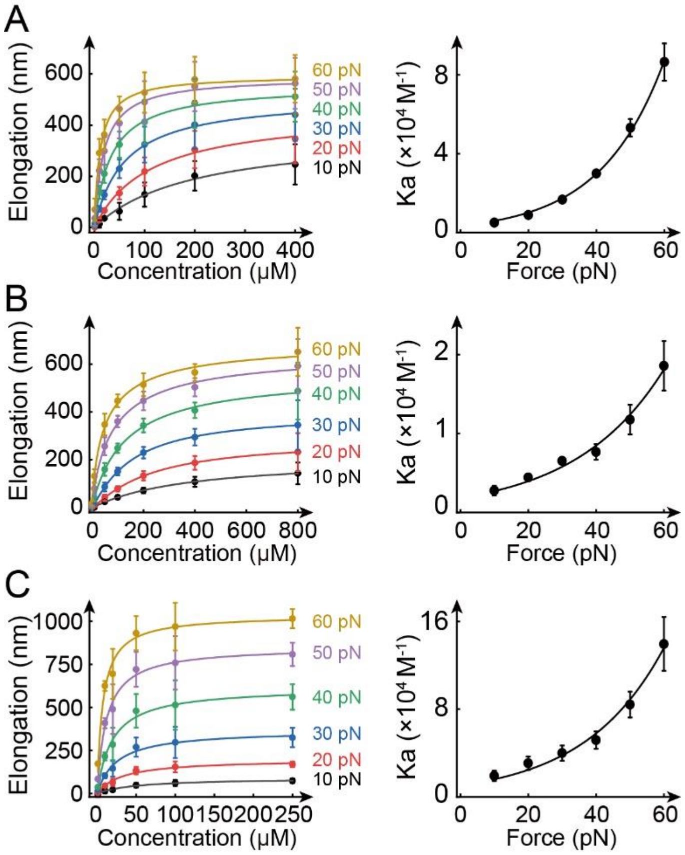

**Figure S4.** Titration curves fit using Hill equation and binding constant dependence on force of compound **2** (A), compound **3** (B) and compound **4** (C).

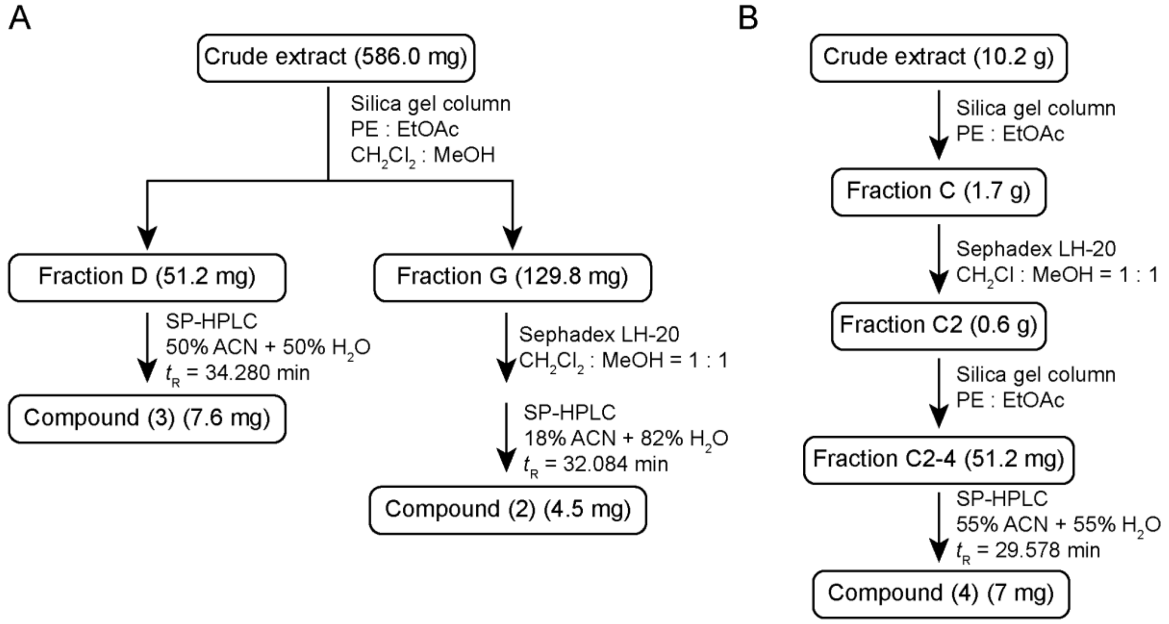

**Figure S5.** Purification of DNA intercalators from bacterium *Streptomyces tanashiensis* (A) and fungi *Talaromyces funiculosus* (B).

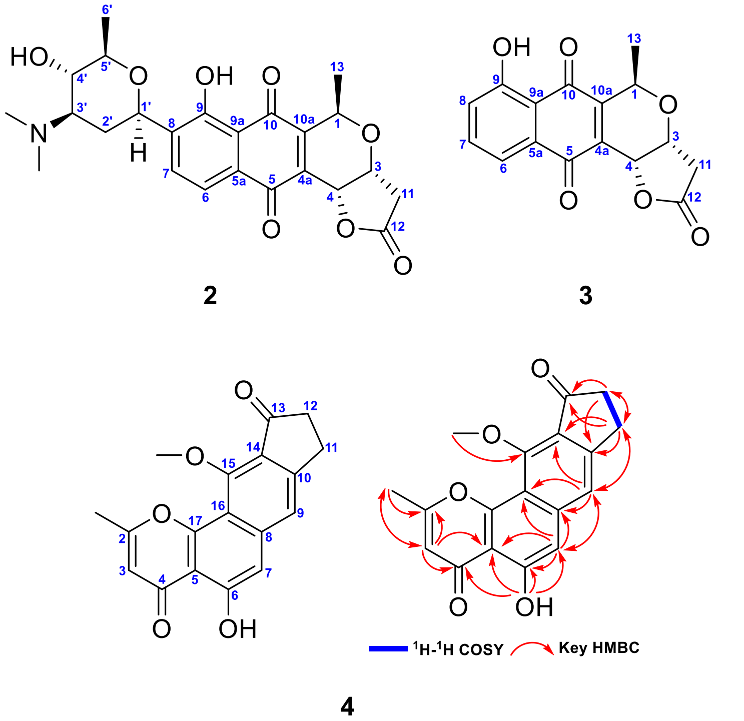

**Figure S6.** Chemical structures of compounds **2-4**, and $^{1}H$-$^{1}H$ COSY and key HMBC correlations of compound **4**.

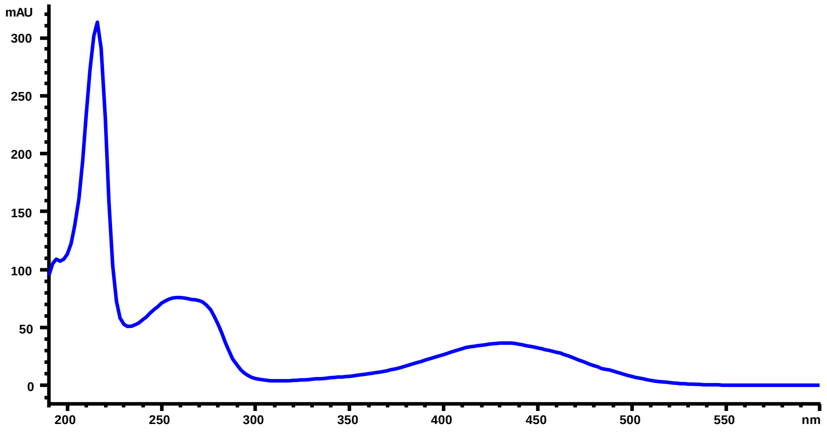

**Figure S7.** UV spectrum of compound **2**.

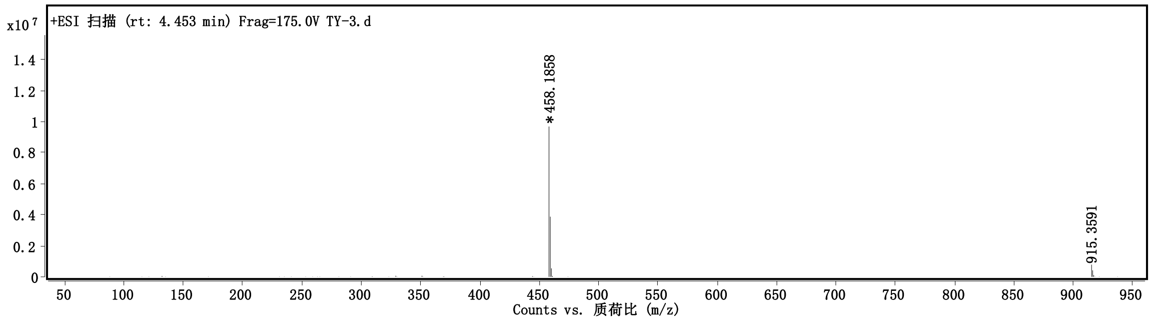

**Figure S8.** HRESIMS spectrum (positive ionization) of compound **2**.

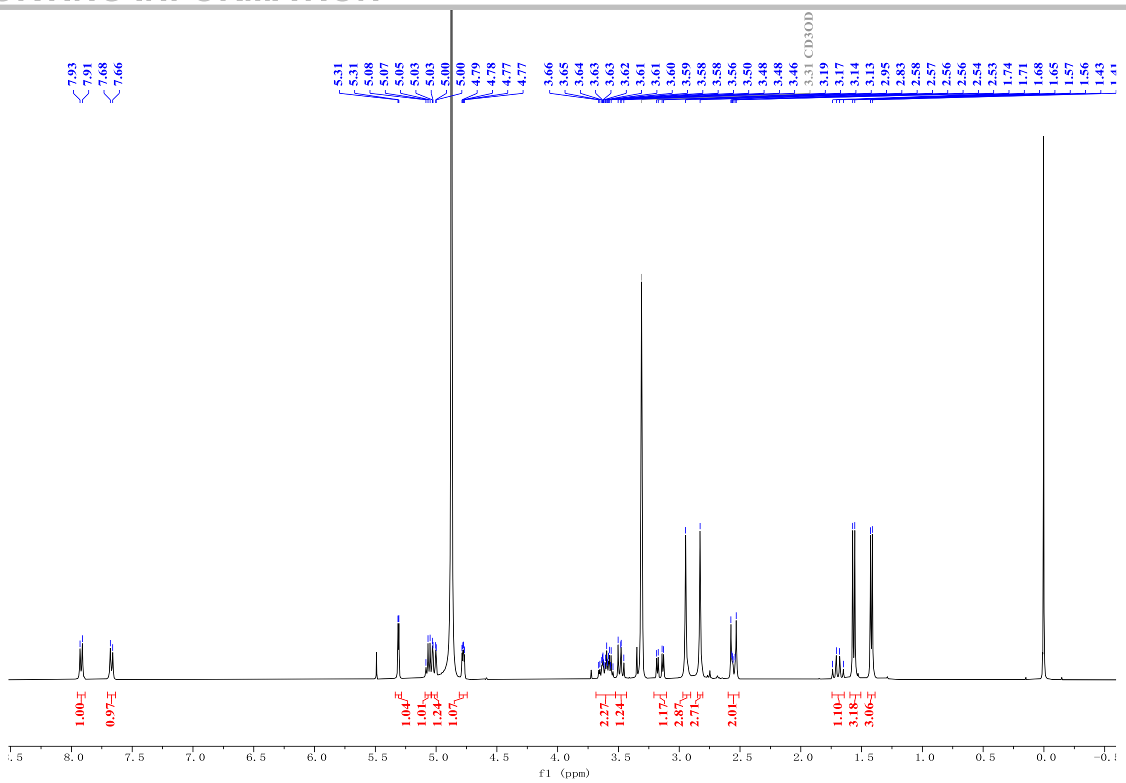

**Figure S9.** $^{1}H$ NMR (400 MHz, $CD_3OD$) spectrum of compound **2**.

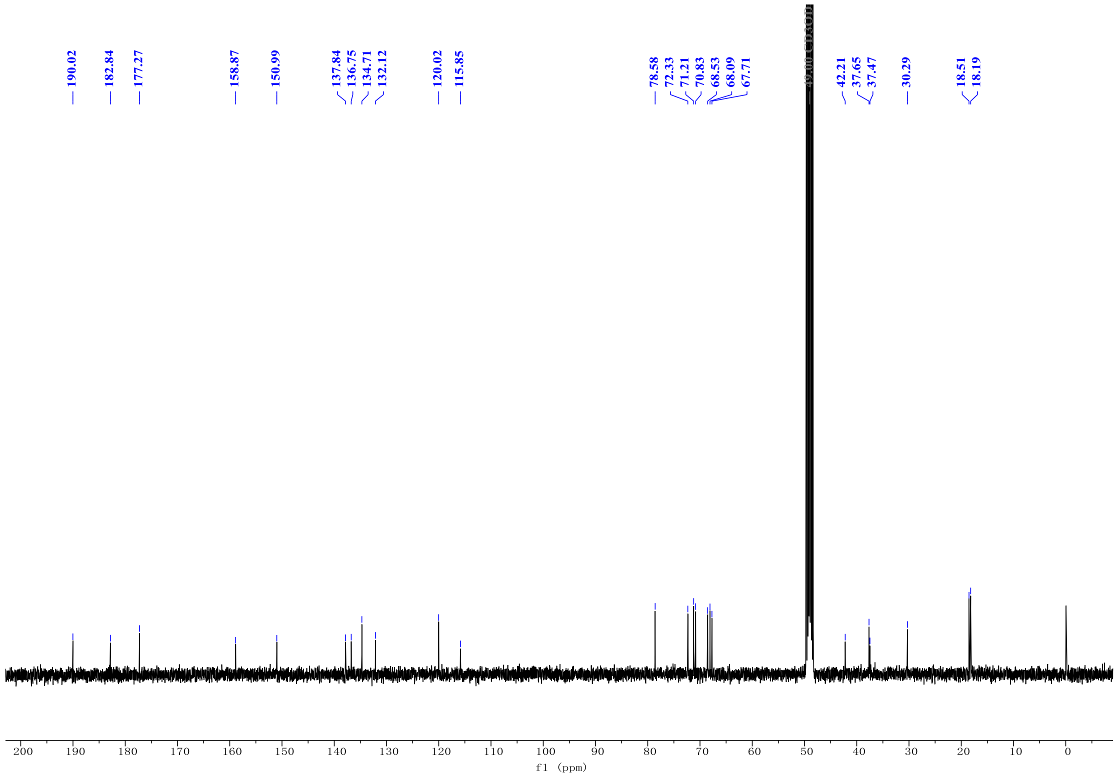

**Figure S10.** $^{13}C$ NMR (100 MHz, $CD_3OD$) spectrum of compound **2**.

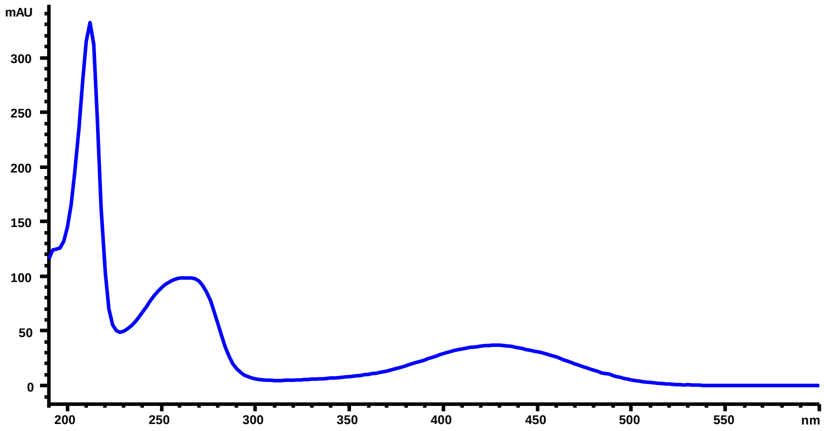

**Figure S11.** UV spectrum of compound **3**.

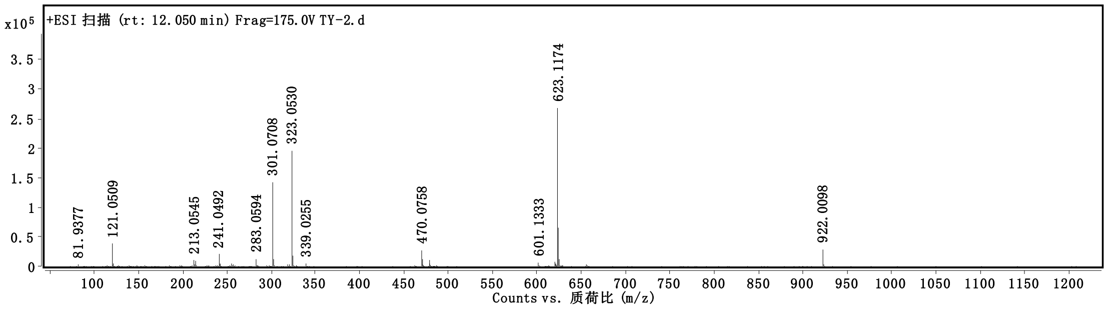

**Figure S12.** HRESIMS spectrum (positive ionization) of compound **3**.

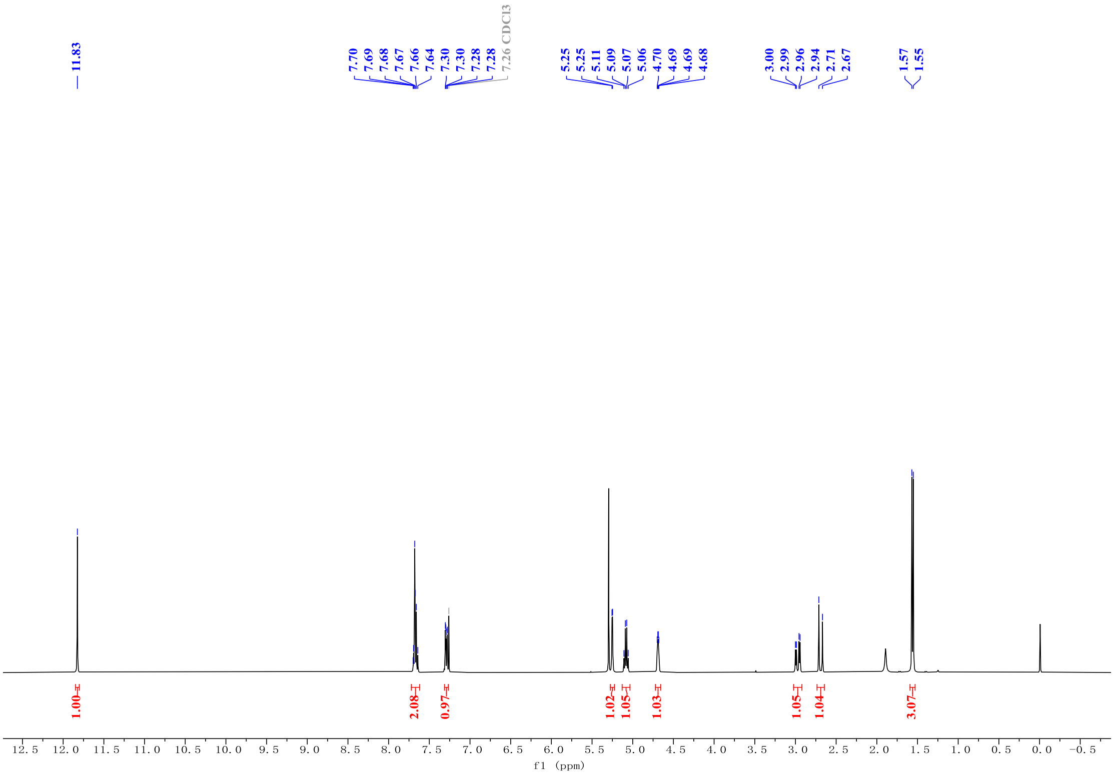

**Figure S13.** $^{1}H$ NMR (400 MHz, $CDCl_3$) spectrum of compound **3**.

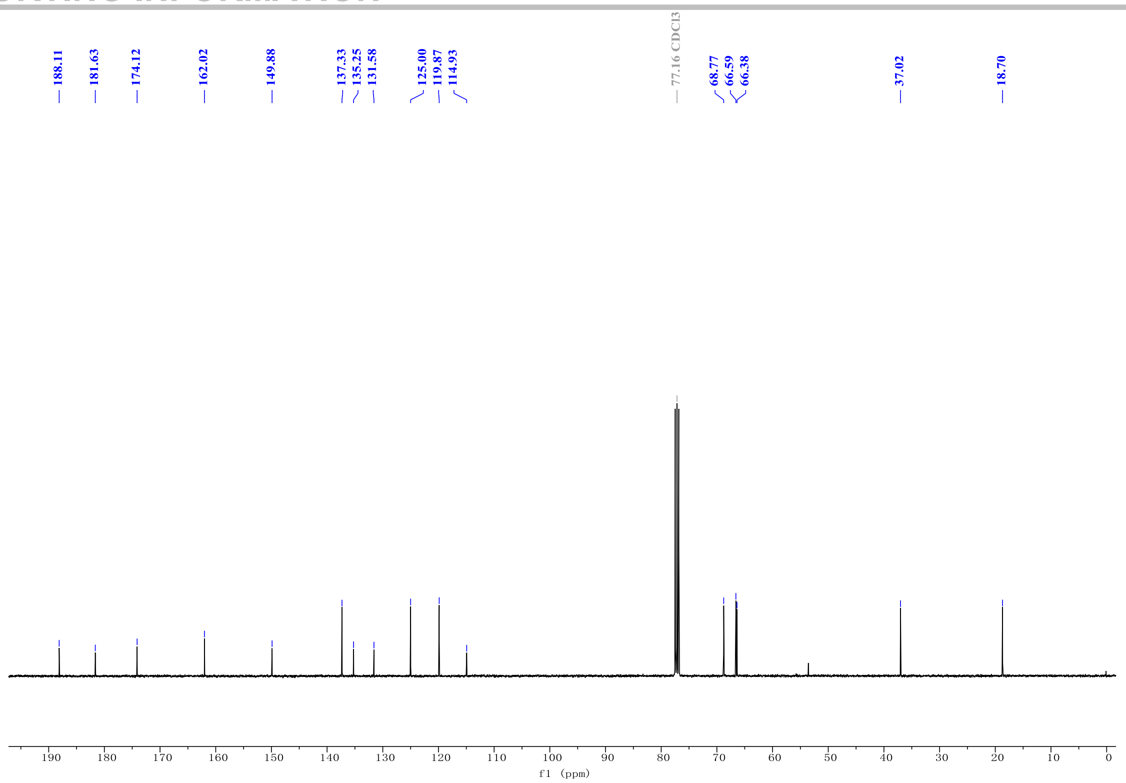

**Figure S14.** $^{13}C$ NMR (100 MHz, $CDCl_3$) spectrum of compound **3**.

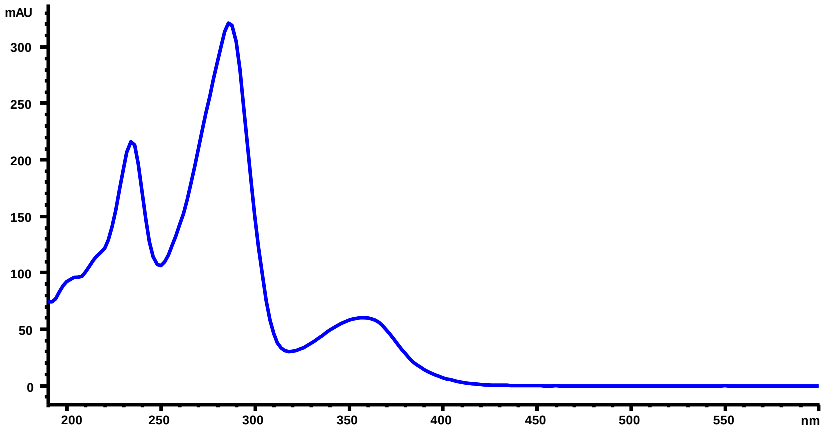

**Figure S15.** UV spectrum of compound **4**.

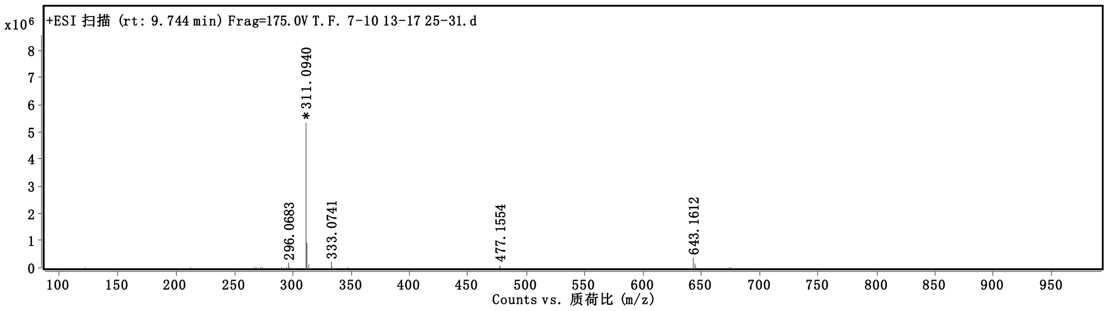

**Figure S16.** HRESIMS spectrum (positive ionization) of compound **4**.

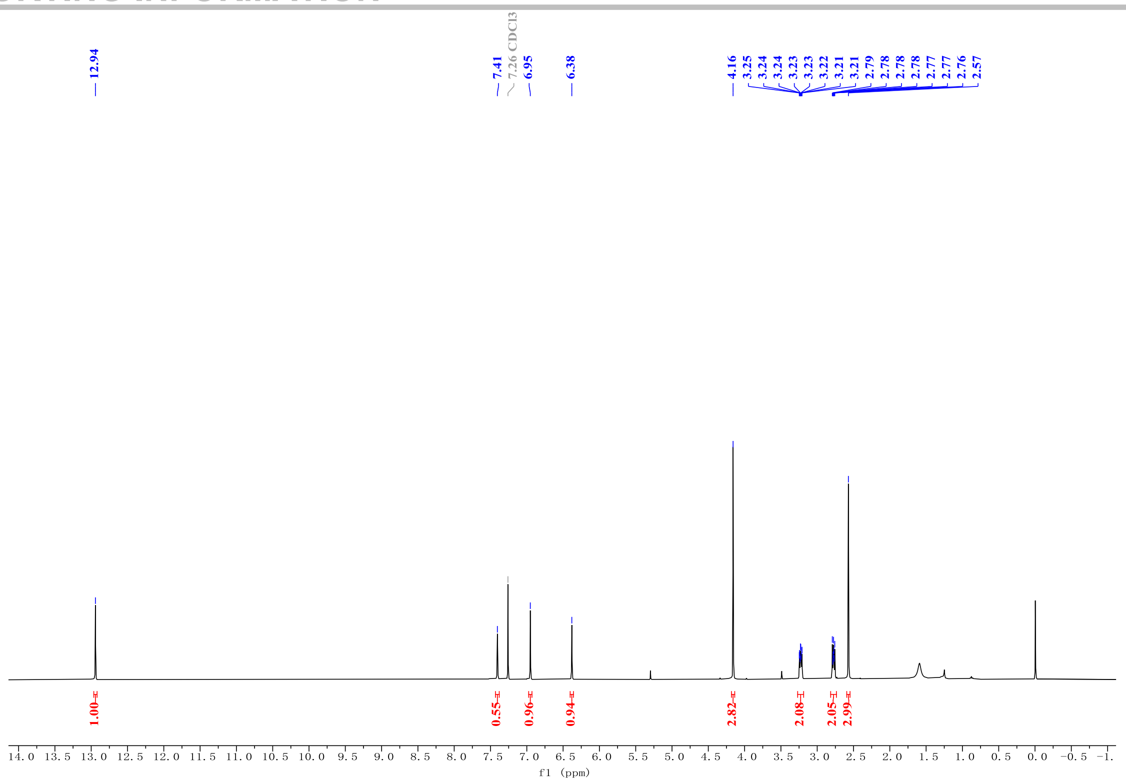

**Figure S17.** $^{1}H$ NMR (400 MHz, $CDCl_3$) spectrum of compound **4**.

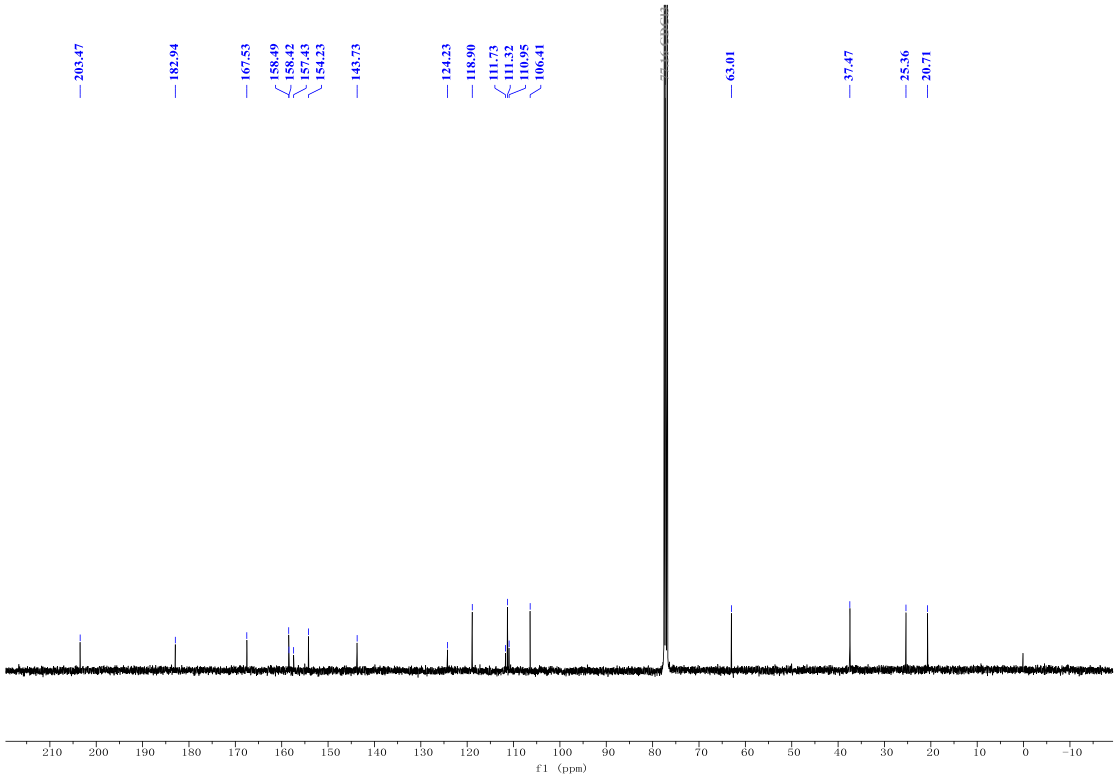

**Figure S18.** $^{13}C$ NMR (100 MHz, $CDCl_3$) spectrum of compound **4**.

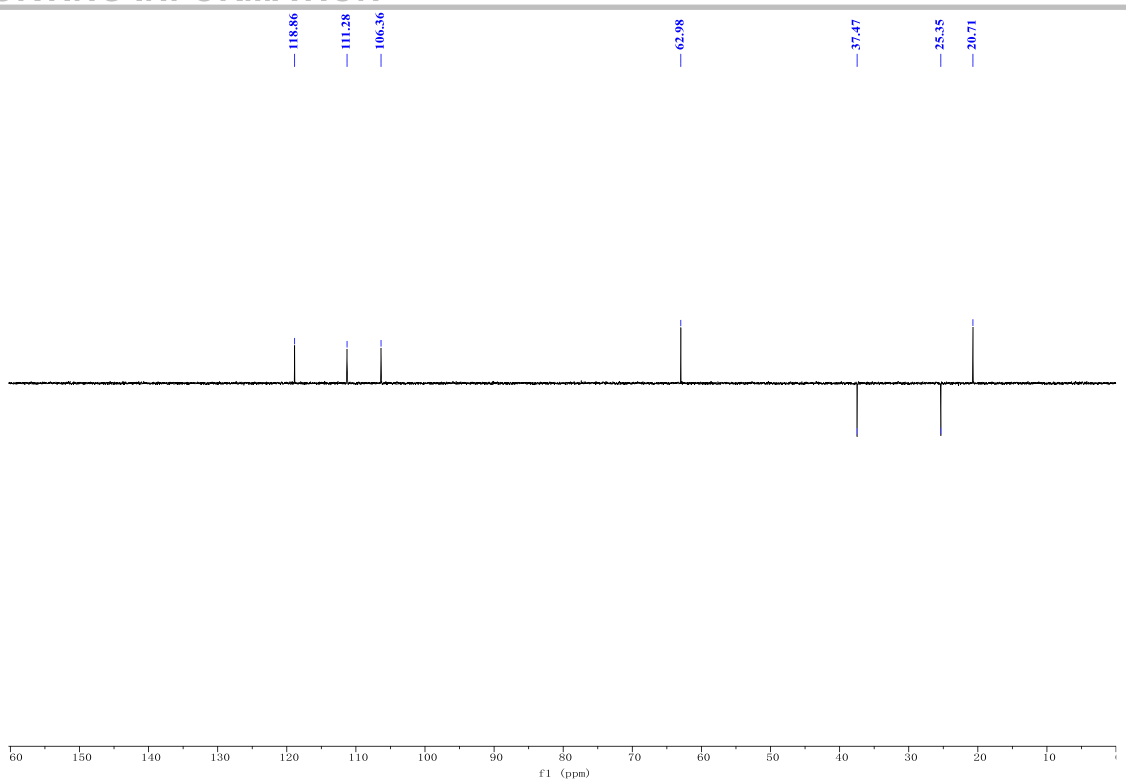

**Figure S19.** DEPT-135° spectrum of compound **4**.

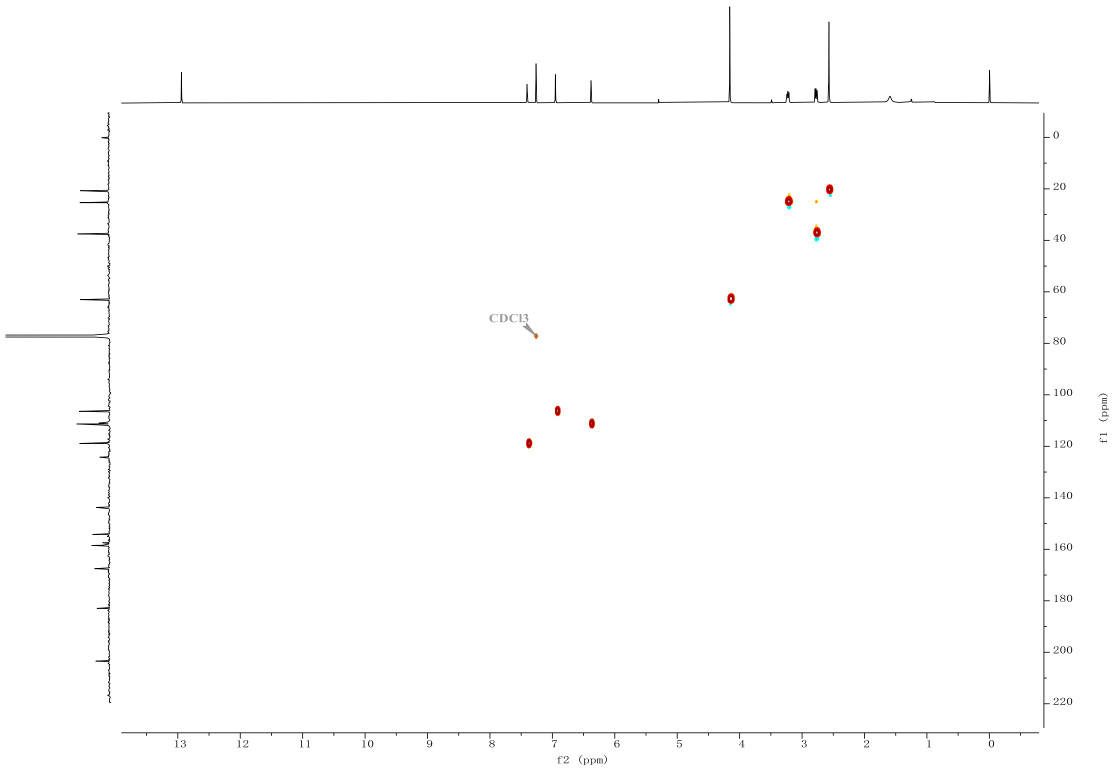

**Figure S20.** HSQC spectrum for compound **4**.

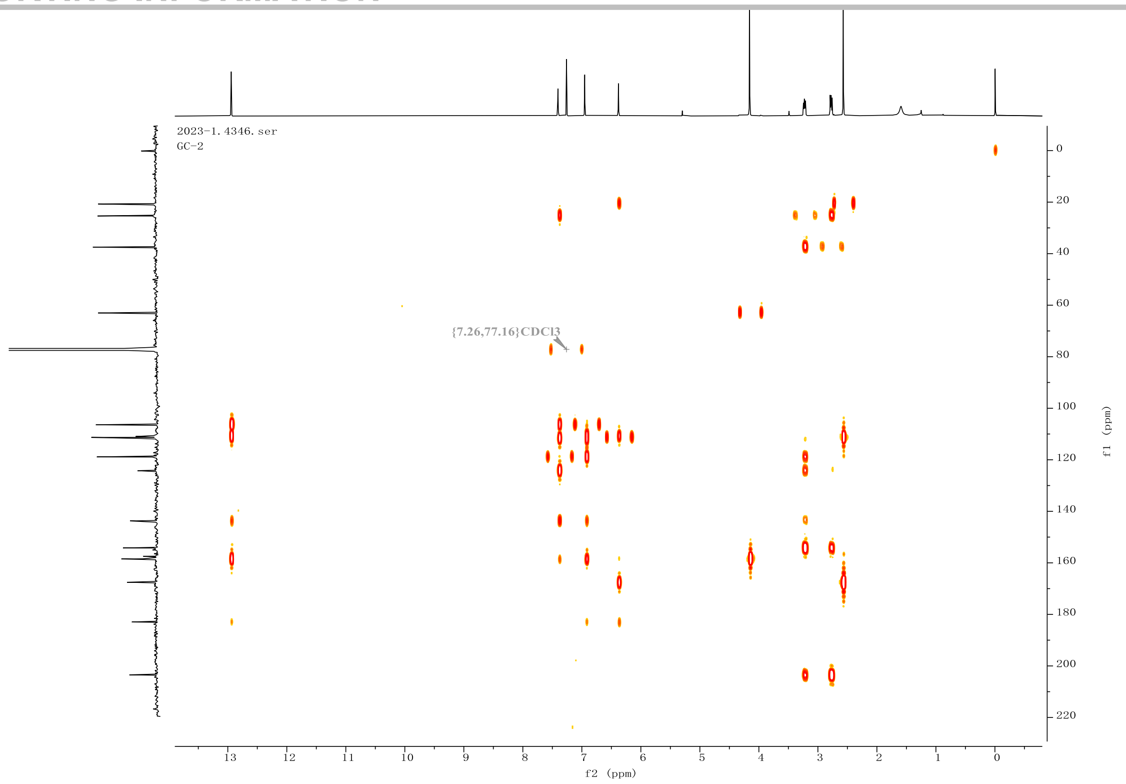

**Figure S21.** HMBC spectrum for compound **4**.

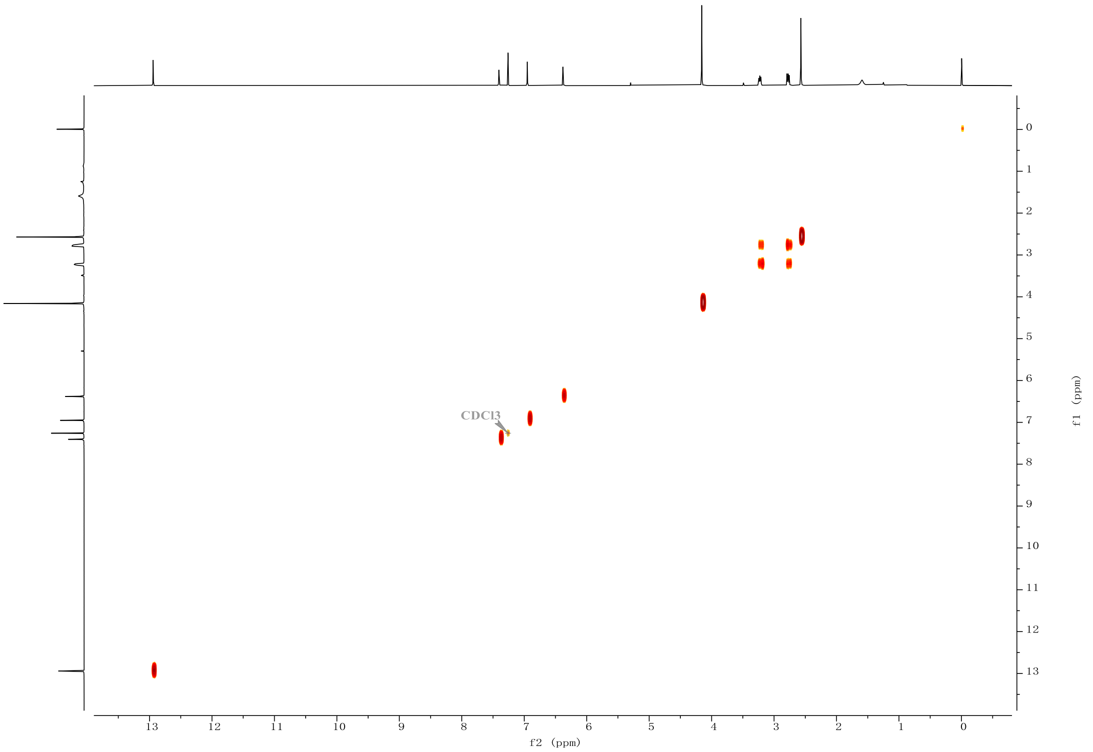

**Figure S22.** $^{1}H$-$^{1}H$ COSY spectrum for compound **4**.
